# Cell viscosity influences hematogenous dissemination and metastatic extravasation of tumor cells

**DOI:** 10.1101/2024.03.28.587171

**Authors:** Valentin Gensbittel, Gautier Follain, Louis Bochler, Klemens Uhlmann, Olivier Lefèbvre, Annabel Larnicol, Sébastien Harlepp, Ruchi Goswami, Salvatore Girardo, Vincent Hyenne, Vincent Mittelheisser, Martin Kräter, Daniel Balzani, Jochen Guck, Naël Osmani, Jacky G. Goetz

**Affiliations:** Tumor Biomechanics, INSERM UMR_S1109, Strasbourg, France; Université de Strasbourg, Strasbourg, France; Fédération de Médecine Translationnelle de Strasbourg (FMTS), Strasbourg, France; Équipe Labellisée Ligue Contre le Cancer; Chair of Continuum Mechanics, Ruhr-Universität Bochum, Universitätsstraße 150, 44801, Bochum, Germany; Max Planck Institute for the Science of Light & Max-Planck-Zentrum für Physik und Medizin, Erlangen, Germany; Department of Physics, Friedrich-Alexander Universität Erlangen-Nürnberg (FAU), Erlangen, Germany

**Keywords:** metastasis, viscosity, elasticity, extravasation, hydrogel beads

## Abstract

Metastases arise from a multi-step process during which tumor cells change their mechanics in response to microenvironmental cues. While such mechanical adaptability could influence metastatic success, how tumor cell mechanics directly impacts intravascular behavior of circulating tumor cells (CTCs) remains poorly understood. In the present study, we demonstrate how the deformability of CTCs affects hematogenous dissemination and identify the mechanical profiles that favor metastatic extravasation. Combining intravital microscopy with CTC-mimicking elastic beads and mechanically-tuned tumor cells, we demonstrate that the inherent properties of circulating objects dictate their ability to enter constraining vessels. We identify cellular viscosity as the key property that governs CTC circulation and arrest patterns. We further demonstrate that cellular viscosity is required for efficient extravasation and find that properties that favor extravasation and subsequent metastatic outgrowth can be opposite. Altogether, we identify CTC viscosity as a key biomechanical parameter that shapes several steps of metastasis.

## Introduction

Metastatic development represents the most dangerous aspect of cancer disease (Steeg, 2016). Life-threatening metastases result from a complex multi-step process, called the metastatic cascade, that includes a step of dissemination via body fluids that allows tumor cells to reach distant organs (Follain et al., 2020). Biomechanical forces challenge tumor cells throughout the entirety of the metastatic process. Known examples include increasing compressive stress at the primary tumor site (Stylianopoulos et al., 2012) or the hemodynamic and collision forces that often threaten to destroy circulating tumor cells (CTCs) (Regmi et al., 2017 ; Wirtz et al., 2011). As such, the mechanical properties of tumor cells likely represent a critical feature in their ability to cope with such mechanical forces throughout the metastatic cascade (Matthews et al., 2020 ; Moose et al., 2020).

While it is commonly accepted that mechanical softness of tumor cells scales with their metastatic potential (Guck et al., 2005 ; Swaminathan et al., 2011 ; Alibert et al., 2017), we recently speculated that the impact of tumor cell deformability on the success of each step might vary at different stages of the metastatic process (Gensbittel et al., 2021). Surprisingly, whether inherent mechanics of CTCs directly impacts hematogenous dissemination remains poorly documented. Although we and others have demonstrated that CTC adhesion potential may control their arrest in non-constraining vessels (Gassmann et al., 2009 ; Osmani et al., 2019), CTC shuttling is drastically reduced when reaching vascular branch points or tiny capillaries that represent highly probable arrest sites for CTCs (Kienast et al., 2010 ; Follain et al., 2018 ; Paul et al., 2019). As such, the ability of CTCs to undergo deformation and accommodate to constraining vessels likely plays an important role in the circulation routes that they take and the vascular sites where they arrest (Tietze et al., 2019).

Some extravasation mechanisms are notoriously dependent on the mechanical features of the extravasating cells. Tumor cells with softer mechanical profiles are shown to be more efficient at performing diapedesis *in vitro* (Chen et al., 2016), during which they can further soften, including their nucleus (Cao et al., 2016 ; Roberts et al., 2021). However, we and other have shown that *in vivo,* tumor cells also exit the bloodstream via endothelial remodeling-driven extravasation (Allen et al., 2019 ; Follain et al., 2018 ; Karreman et al., 2023), a process in which arrested tumor cells are actively extracted from blood vessels by endothelial cells. Whether tumor cell mechanics is involved in this alternate extravasation mechanism remains unknown. As a result, our understanding of the role of cell mechanics in extravasation remains incomplete as the different extravasation mechanisms that have been described might have different mechanical requirements for tumor cells.

In this work, we take advantage of two *in vivo* models and advanced biophysical tools to broaden our understanding of biomechanics in metastasis. We speculated that, and tested whether the inherent mechanics of CTCs profoundly impact their circulation, arrest and extravasation potential in constraining vessels. We exploited elastic cell-mimicking hydrogel (polyacrylamide) beads in combination with the fine-tuning of the viscoelasticity of CTCs to demonstrate, for the first time, that it is the cells’ viscosity and not their elasticity that determines whether they can efficiently enter into, and then arrest in constraining vasculature. We show that high-viscosity tumor cells efficiently extravasate via intraluminal endothelial remodeling in experimental metastasis assays performed in two animal models. Yet, although high viscosity favors extravasation, it does not necessarily translate into increased subsequent metastatic outgrowth, further highlighting the need of tumor cells for mechanical adaptation of their viscoelastic profiles. Altogether, we identify CTC viscosity as a key biomechanical parameter that shapes several intravascular steps of the metastatic cascade (i.e., circulation, arrest and extravasation).

## Results

### Small size and low elasticity of circulating objects facilitate lodging in constraining blood vessels

We and others have demonstrated that CTCs exploit adhesion to efficiently and stably arrest in vascular regions with permissive hemodynamics (Gassmann et al., 2009 ; Follain et al., 2018 ; Osmani et al., 2019). Yet, CTCs often face constraining vascular architectures that force them to stop (Kienast et al., 2010 ; Follain et al., 2018 ; Headley et al., 2016). Thus, we first sought to demonstrate that the mechanics of circulating objects (cell-like beads or tumor cells) also dictates their circulation and arrest patterns independently of their biochemical properties. We exploited the zebrafish embryo, and its stereotyped vasculature, as a means to interrogate spatio-temporal patterns of arrest as a function of the mechanics of the circulating objects. The caudal plexus of the zebrafish embryo offers an ideal window to stereotypical vascular topology with small-size vessels (intersegmental vessels (ISVs) and caudal plexus capillaries (CPCs)) and large vessels (dorsal aorta (DA), arterio-venous junction (AVJ), caudal vein (CV)) offering multiple routes for circulating objects (Figures 1A, S1A). We used cell-like beads (i.e., elastic polyacrylamide beads, (Girardo et al., 2018)), with no adhesive capacity and whose diameters and elasticities were tuned (Figures 1A, S1B), to probe circulation and arrest patterns shortly upon injection in the bloodstream (5 min post-injection recording to document the majority of the arrest events) (Figure 1A). We used an in-house heat-mapping workflow to highlight hotspots of arrest of small, large, soft and stiff circulating beads (Figure 1B). We found that both reduced diameter (19 µm vs 11 µm) and reduced elasticity (2.0 kPa vs 0.3 kPa) led to changes in proportions of beads arresting in the AVJ, the CV, the ISVs and the CPCs (Figures 1C, S1C-D), all-in-all resulting in significant increases of beads managing to enter and arrest in small constraining vessels (ISVs + CPCs) when properties of size or elasticity are reduced (Figure 1D). Interestingly, when comparing multiple tumor cell lines displaying significant differences in cell diameter, we also observed a significant correlation between their size and their capacity to enter and arrest in small vessels, mirroring our bead observations (Figures S1E-G).

**Figure 1:**
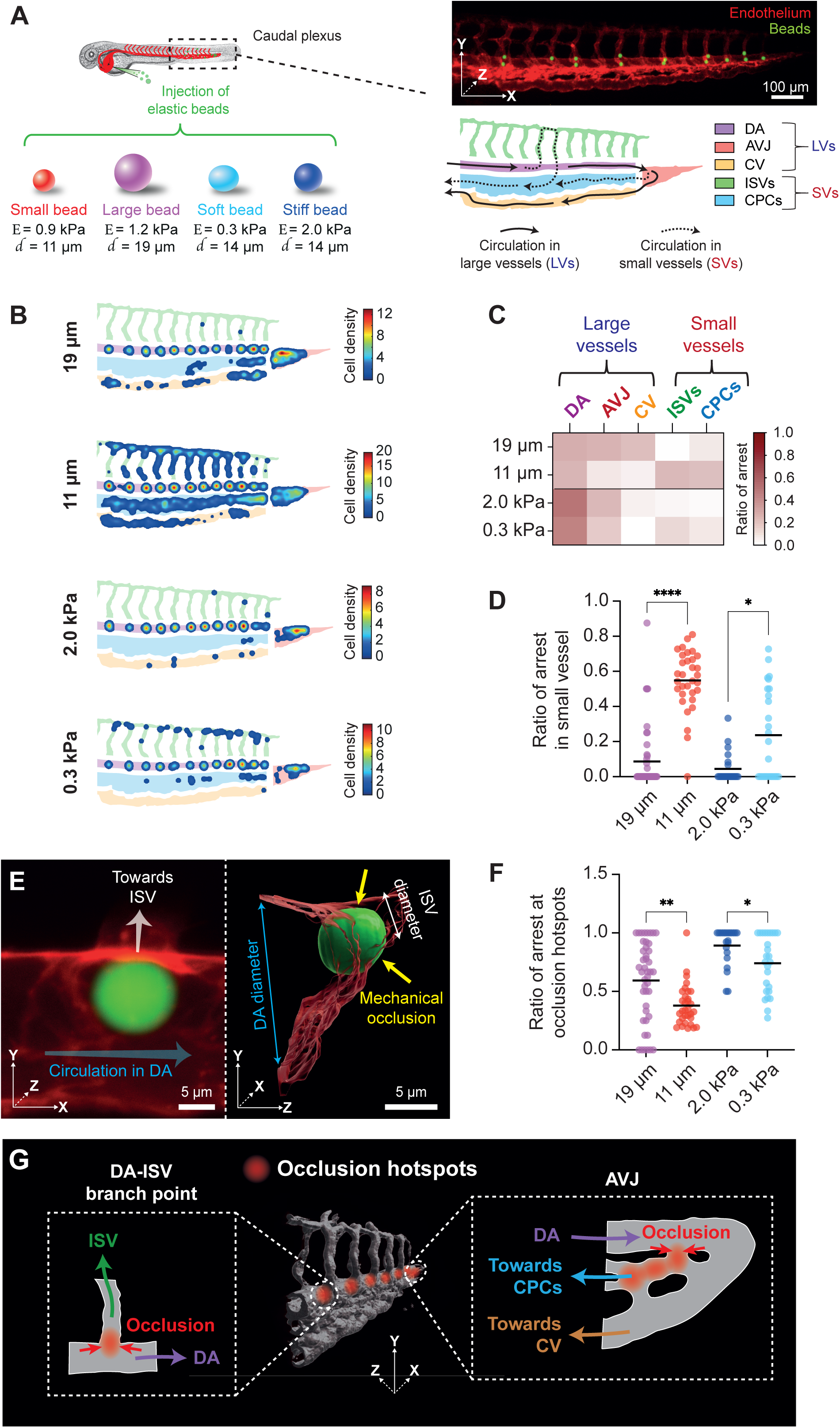
Small size and low elasticity of circulating objects facilitate lodging in constraining blood vessels. (A) Bead populations injected in the duct of Cuvier of the ZF embryo model for imaging their circulation and arrest events in the vascular subregions that constitute the caudal plexus, Stereomicroscope image of arrested beads (green) in the vasculature of the caudal plexus of a ZF embryo (red) 5 minutes post-injection and map breaking down the caudal plexus (Dorsal Aorta (DA) ; Arterio-Venous Junction (AVJ) ; Caudal Vein (CV) ; Intersegmental Vessels (ISVs) ; Caudal Plexus Capillaries (CPCs)) of the ZF embryo. (B) Heatmaps displaying hotspots of arrest 5 mpi of large (19 µm), small (11 µm), stiff (2.0 kPa) and soft (0.3 kPa) polyacrylamide elastic beads in the ZF embryo caudal plexus (N = 2, n (embryos) = 44, 32, 22 and 26 for large, small, stiff and soft beads respectively). (C) Data heatmap of ratios of bead arrest occurring in the vascular subregions of the caudal plexus of the ZF embryo 5 minutes post-injection. (D) Quantification of ratio of arrest occurring in small vessels (ISVs + CPCs) of the ZF embryo caudal plexus 5 minutes post-injection of large (19 µm), small (11 µm), stiff (2.0 kPa) and soft (0.3 kPa) polyacrylamide elastic beads. (N = 2, n (embryos) = 44, 32, 22 and 26 for large, small, stiff and soft beads respectively). (Mann-Whitney, p-values = <0.0001 (Large vs Small) ; 0.0103 (Stiff vs Soft)). (E) 3D renders of a bead occluding the entrance of an ISV at a DA-ISV vascular branch point. (F) Quantification of ratio of arrest occurring at hotspots of mechanical occlusion in the ZF embryo caudal plexus 5 minutes post-injection of large (19 µm), small (11 µm), stiff (2.0 kPa) and soft (0.3 kPa) polyacrylamide elastic beads. (N = 2, n (embryos) = 44, 32, 22 and 26 for large, small, stiff and soft beads respectively). (Mann-Whitney, p-values = 0.0017 (Large vs Small) ; 0.0329 (Stiff vs Soft)). (G) Graphical summary of main occlusion sites within the ZF embryo caudal plexus vasculature.

Mechanical occlusion was the major driver of the arrest of non-adherent circulating beads as exemplified with 3D reconstruction of arrested beads imaged with high-resolution confocal imaging (Figure 1E). We found that properties of reduced size 19 µm vs 11 µm) and reduced elasticity (2.0 kPa vs 0.3 kPa) both led to significant decreases in proportions of beads arrested at the major occlusion sites (Figure 1F) that we identified at the vascular branch points (for example, at DA-ISV connections or at sites where the DA gets subdivided into the CV and the CPCs) (Figure 1G). Altogether, these results suggest that size, but also elasticity, of circulating objects dictate their capacity to enter and stop in a mechanically-constraining vascular system often encountered in organs of metastasis. Reduced diameter or increased compliance allow them to escape major occlusion sites and thereby favor entry and arrest in small blood vessels.

### CTCs use viscous deformation to enter small-sized vessels

We next sought to characterize how objects trapped within the vasculature cope with the constraints of the intravascular environment. To do so and to probe the stresses at play at occlusion sites, we exploited inert tumor cell-like polyacrylamide beads, with a defined elasticity (0.8 kPa) (Figure 2A), as cell-like force sensors (Träber et al., 2019) and computed the volumetric mean of pressure values upon high-resolution 3D imaging at their site of vascular arrest (Figures 2B-C, S2A) and with respect to their position (Figure S2B). Unexpectedly, we observed no difference in volumetric mean of pressure between beads arrested in various vascular environments (Figure S2C). No difference was observed between beads arrested in large (LVs) vs small vessels (SVs) (Figure 2D) or between beads arrested at main occlusion sites previously identified vs other sites (Figure S2D). Consistently, our shape factor analysis of the bead geometry of these same beads did not highlight any significant differences in bead volume (Figure S2E), in bead elongation (Figure 2E) or in bead flatness (Figure S2F) between beads located in small vs large vessels. This shows that elastic beads are poorly constrained once arrested and suggests either that the endothelial wall of vessels reshapes in response to their presence or, alternatively, that the amount of stress is not sufficient to significantly deform these cell-like elastic objects that remain occluded. Aiming at further exploring which cell mechanical phenotypes dictate successful entry, via deformation, in small-sized vessels, we applied the same shape factor analysis to arrested tumor cells (Figures 2F, S2G). When applying high-resolution 3D visualization of tumor cell geometry (Figure 2G), we found the average cell volume of arrested tumor cells to be significantly smaller for cells arrested in small vessels compared to ones arrested in large vessels (Figure 2H). Both cell elongation (Figure 2I) and cell flatness (Figure S2H) were significantly higher for cells arrested in small vessels, suggesting that contrary to elastic beads, viscoelastic tumor cells do significantly change their shape to adapt to the constraining vascular architecture (Figure S2I-K).

**Figure 2:**
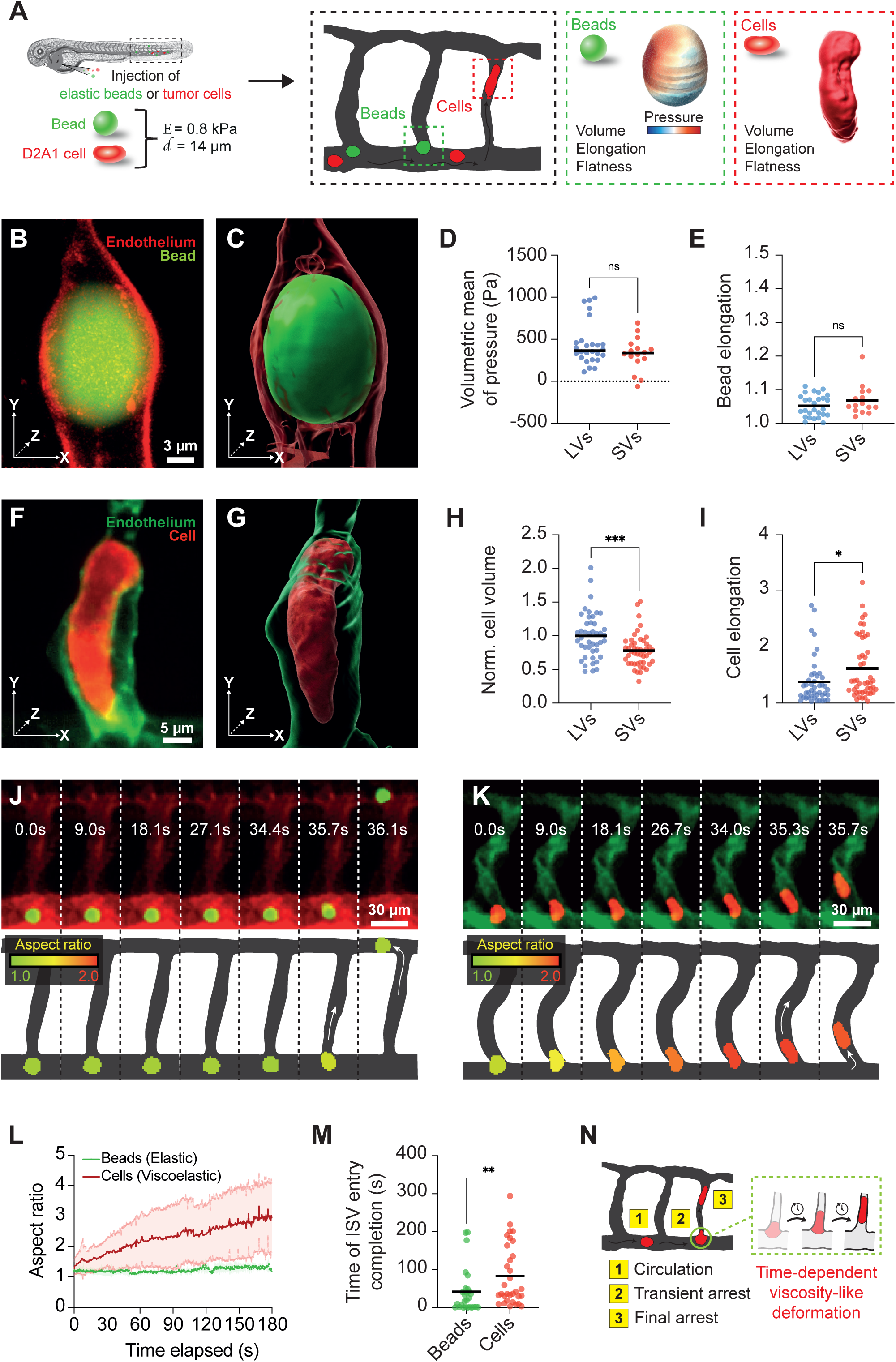
CTCs use viscous deformation to enter small-sized vessels. (A) Workflow for bead/cell deformation analysis and computation of mechanical stress experienced by beads in the intravascular environment of the ZF embryo. (B) Z-projection of confocal stack of arrested “D2A1-like” bead in an ISV. (C) 3D renders of (B). (D) Quantification of volumetric mean of pressure sustained by arrested beads in large (LVs) vs small (SVs) vessels. (N = 6, n (beads) = 26 and 16 for LVs and SVs respectively) (Mann-Whitney, p-value = 0.2978). (E) Quantification of elongation of beads arrested in LVs and SVs (N = 6, n (beads) = 26 and 16 for LVs and SVs respectively) (Mann-Whitney, p-value = 0.254). (F) Z-projection of confocal stack of arrested D2A1 tumor cell in an ISV. (G) 3D renders of (F). (H) Quantification of volume of D2A1 tumor cells arrested in LVs and SVs. (N = 2, n (cells) = 44 and 47 for LVs and SVs respectively) (Mann-Whitney, p-value = 0.0005). (I) Quantification of elongation of D2A1 tumor cells arrested in LVs and SVs (N = 2, n = 44 and 47 for LVs and SVs respectively) (Mann-Whitney, p-value = 0.0128). (J) Stereomicroscope time lapse and aspect ratio tracking of polyacrylamide elastic bead overcoming an occlusion site at a DA-ISV vascular branch point. (K) Stereomicroscope time lapse and aspect ratio tracking of D2A1 tumor cell overcoming an occlusion site at a DA-ISV vascular branch point. (L) Aspect ratio as a function of time spent at ISV entrance for polyacrylamide elastic beads and D2A1 tumor cells (N = 6 and 2 for beads and cells respectively, n (events of arrival, arrest and entry at DA-ISV connections) = 26 and 32 for beads and cells respectively). (M) Time spent at ISV entrance before completing entry for elastic beads and D2A1 tumor cells. (N = 6 and 2 for beads and cells respectively, n (events of arrival, arrest and entry at DA-ISV connections) = 26 and 32 for beads and cells respectively) (Mann-Whitney, p-value = 0.0095). (N) Graphical summary of viscous-like deformation of cells overcoming occlusion sites and entering constraining vessels.

Based on this observation that elastic beads struggle to deform their shape compared to viscoelastic cells despite being set at a comparable elasticity level, we next selected our most compliant 0.3 kPa bead population to be compared to cells when tracking their aspect ratio as they go through the process of entering in small constraining ISVs (Figure 2J-K). During the transient arrest at these occlusion sites, beads appeared to keep a constant aspect ratio of spherical nature, while tumor cells progressively deformed and increased their aspect ratio with time on a minute timescale (Figure 2J-L). We normalized both the change in aspect ratio and the amount of time spent at the ISV entrance before completing entry for both tumor cells and beads (Figure S2L). This allowed us to confirm, on one hand, that tumor cells did indeed progressively and drastically deform their shape, in a viscous-like manner, over the entire duration of time that they spent arrested at these occlusion sites before completing entry. On the other hand, this allowed us to reveal that the beads that previously appeared to display stable aspect ratios at the occlusion sites actually underwent deformation as well, only this deformation was very sudden and transient and persisted only in the fractions of seconds that preceded their entry into ISVs (Figure S2L), in line with their elastic nature. Interestingly, although beads had a low elasticity (0.3 kPa) compared to that of cells (≈ 0.8 kPa), the extent to which they deformed remained minor compared to cells (Figure S2M). As a result of the time-dependent nature of the deformation for viscoelastic cells, the average time that was required for them to overcome the occlusion sites and complete entry in constraining ISVs was significantly increased compared to elastic beads (Figure 2M). Altogether, these results demonstrate that viscoelastic tumor cells exploit their viscosity to overcome the mechanical challenges occurring at occlusion sites and progressively creep into smaller vessels across longer timescales reminiscent of viscous behaviors. (Figure 2N).

### Cell viscosity dictates arrest locations of circulation tumor cells

Given that we identified viscous-like deformation in CTCs overcoming occlusion sites but yet also demonstrated that tuning elasticity in an elastic system (beads) impacted the ability to enter constraining vessels, we set out to better dissect the respective contributions of elasticity and viscosity in a fully viscoelastic cell mechanics model. In order to modify viscoelastic properties of tumor cells, we targeted several structures controlling overall cell mechanics (nucleus : lamin A/C ; intermediate filaments : vimentin ; actomyosin network: non-muscle myosin IIA ; plasma membrane : caveolin-1) using RNAi silencing (Figure S3A) and measured the resulting viscoelasticity using a micropipette aspiration assay which mimics, in space and time, the *in vivo* situation where CTCs escape occlusion sites and creep into SVs (Figures 3A, S3B). While all RNAi treatments significantly decreased the elasticity of tumor cells (Figure 3B), depleting MYH9 had no effect on cell viscosity (Figure 3C). We exploited such results to distinguish cells where neither elasticity nor viscosity were altered (siCTRL), cells where only elasticity was decreased (siMYH9) and cells in which both elasticity and viscosity were significantly decreased (siVIM, siCAV1, siLMNA) for performing *in vivo* experiments in the zebrafish embryo (Figure 3D). We recorded the arrest events for 5 minutes post-injection (Figure 3E), applied our heatmapping workflow (Figures 3F, S3C) and quantified the arrest events in all vascular subregions of the ZF embryo caudal plexus (Figure 3G). When quantifying arrest in small vessels, we observed that low viscosity tumor cells (siVIM, siCAV1, siLMNA) better entered and arrested in small vessels (Figure 3H). Interestingly, CTCs with low elasticity but unchanged viscosity levels (siMYH9) behaved similar to control CTCs (siCTRL). Concomitantly, we observed the opposite trend when quantifying the arrest events at major occlusion hotspots, with low viscosity tumor cells (siVIM, siCAV1, siLMNA) displaying reduced proportions of cells stuck at these arrest sites compared to high viscosity cells (siCTRL, siMYH9) (Figure 3H). These results suggest that cell viscosity is a key determinant of CTCs getting past occlusion sites and entering into small constraining vessels (Figure 3J). Indeed, when plotting the ratios of arrest in SVs as a function of CTC viscosity, we observed that cellular viscosity was strongly correlated with their arrest in SVs (Figure 3K). We then successfully increased viscosity, and not elasticity, of tumor cells by over-expressing vimentin (Figures S3D-E). As expected, such opposite mechanical tuning reduced the capacity of CTCs to enter small constraining vessels (Figures S3F-G).

**Figure 3:**
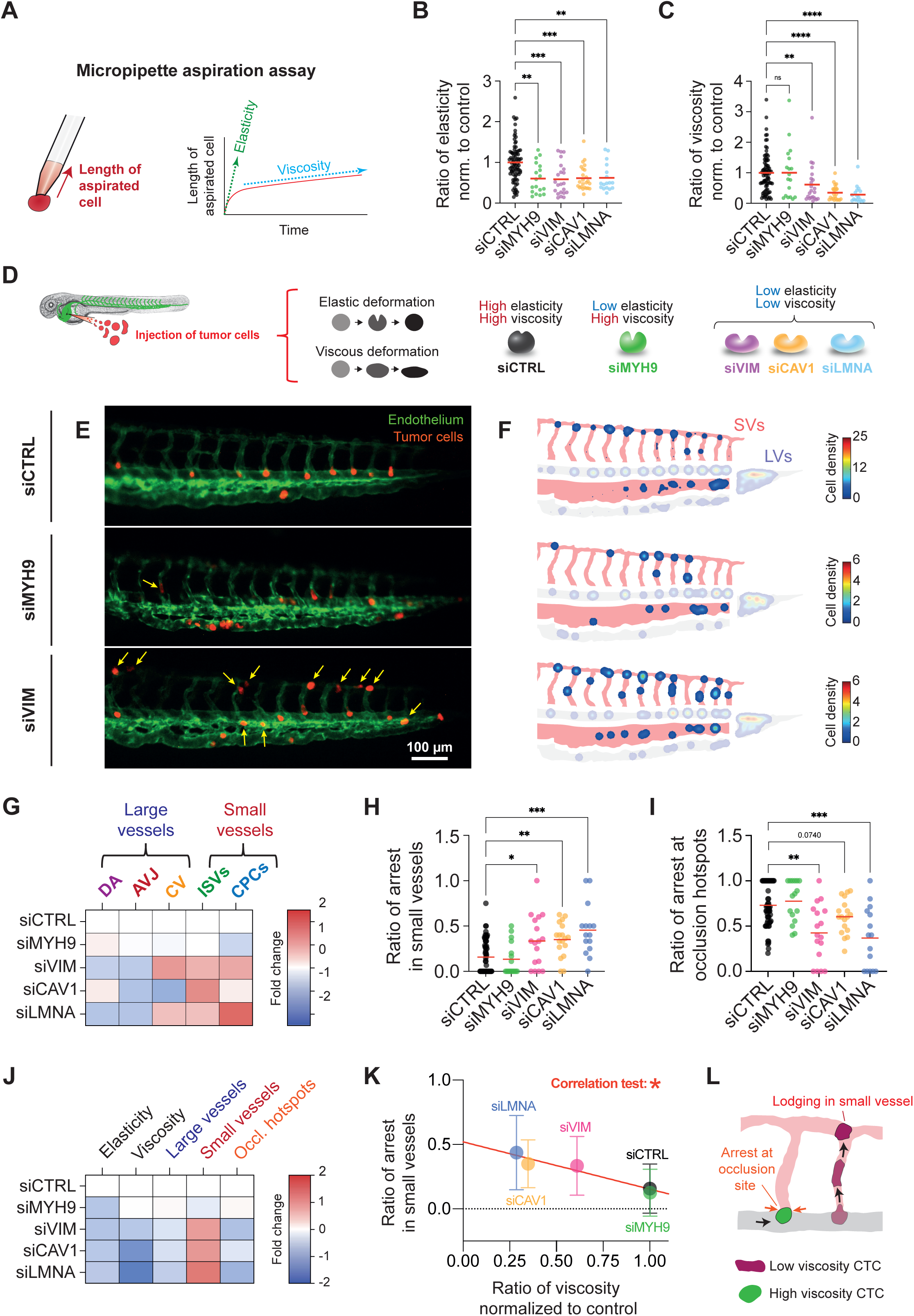
Cell viscosity dictates arrest locations of circulating tumor cells. (A) Micropipette aspiration assay for measuring viscoelastic properties of tumor cells. (B) Ratio of elasticity of siRNA-treated D2A1 tumor cells normalized to control. (N = 11 (siCTRL) ; 3 (siMYH9, siVIM, siCAV1) ; 2 (siLMNA), n (cells) = 82 (siCTRL) ; 18 (siMYH9) ; 22 (siVIM) ; 24 (siCAV1) ; 17 (siLMNA)) (Kruskal-Wallis, p-values = 0,0011 (siCTRL vs siMYH9) ; 0,0005 (siCTRL vs siVIM) ; 0,0009 (siCTRL vs siCAV1) ; 0,0030 (siCTRL vs siLMNA)). (C) Ratio of viscosity of siRNA-treated D2A1 tumor cells normalized to control. (N = 11 (siCTRL) ; 3 (siMYH9, siVIM, siCAV1) ; 2 (siLMNA), n (cells) = 82 (siCTRL) ; 18 (siMYH9) ; 22 (siVIM) ; 24 (siCAV1) ; 17 (siLMNA)) (Kruskal-Wallis, p-values = 0,5585 (siCTRL vs siMYH9) ; 0,0033 (siCTRL vs siVIM) ; <0,0001 (siCTRL vs siCAV1) ; <0,0001 (siCTRL vs siLMNA)). (D) Injection of mechanically-altered tumor cells in the zebrafish embryo. (E) Merged stereomicroscope images of D2A1 tumor cells arrested in the caudal plexus of ZF embryo 5 minutes post-injection. Yellow arrows indicate tumor cells arrested in small vessels (ISVs + CPCs). (F) Heatmaps displaying hotspots of arrest 5 minutes post-injection of siRNA-treated D2A1 tumor cells in the ZF embryo caudal plexus (N = 12 (siCTRL) ; 3 (siMYH9, siVIM), n (embryos) = 59 (siCTRL) ; 16 (siMYH9) ; 17 (siVIM)). (G) Fold change of ratio of arrest occurring in the vascular subregions of the caudal plexus of the ZF embryo 5 minutes post-injection of mechanically-altered tumor cells. (N = 12 (siCTRL) ; 3 (siMYH9, siVIM, siCAV1, siLMNA), n (embryos) = 59 (siCTRL) ; 16 (siMYH9) ; 17 (siVIM) ; 18 (siCAV1) ; 15 (siLMNA)). (H) Quantification of ratio of arrest occurring in small vessels (ISVs + CPCs) of the ZF embryo caudal plexus 5 minutes post-injection of mechanically-altered tumor cells. (N = 12 (siCTRL) ; 3 (siMYH9, siVIM, siCAV1, siLMNA), n (embryos) = 59 (siCTRL) ; 16 (siMYH9) ; 17 (siVIM) ; 18 (siCAV1) ; 15 (siLMNA)) (Kruskal-Wallis, p-values = 0,7125 (siCTRL vs siMYH9) ; 0,0193 (siCTRL vs siVIM) ; 0,0014 (siCTRL vs siCAV1) ; 0,0004 (siCTRL vs siLMNA)). (I) Quantification of ratio of arrest occurring at mechanical occlusion hotspots of the ZF embryo caudal plexus 5 minutes post-injection of mechanically-altered tumor cells. (N = 12 (siCTRL) ; 3 (siMYH9, siVIM, siCAV1, siLMNA), n (embryos) = 59 (siCTRL) ; 16 (siMYH9) ; 17 (siVIM) ; 18 (siCAV1) ; 15 (siLMNA)) (Kruskal-Wallis, p-values = 0,5922 (siCTRL vs siMYH9) ; 0,0011 (siCTRL vs siVIM) ; 0,0740 (siCTRL vs siCAV1) ; 0,0009 (siCTRL vs siLMNA)). (J) Fold changes of ratios of elasticity, viscosity, arrest in small and large vessels and arrest at occlusion hotspots for mechanically-altered tumor cells. (K) Correlation test of ratio of arrest occurring in small vessels (G) and ratio of viscosity (C). (Spearman r, p-value = 0.0167). (L) Graphical summary of the impact of cell viscosity on circulation and arrest patterns of circulating tumor cells.

While the effect did not reach statistical significance, which can in part be explained by the low frequency of such events (ratio approximating 0,18 for CTRL cells), it further increased the statistical significance of the correlation of viscosity and arrest in small vessels (Figures S3H-I). Such correlation is absent with elasticity (Figure S3J). Altogether, these results further confirm that viscosity tunes tumor cells’ ability to overcome occlusion sites and show that modulating cell viscosity has a profound impact on the dissemination and arrest patterns of CTCs (Figure 3L).

### Tumor cell viscosity tunes metastatic extravasation

Metastatic outgrowth requires, in most cases, that CTCs extravasate from the intravascular environment they have reached. Having demonstrated that cellular viscosity controls their ability to reach constraining vessels, we wondered whether it also impacted the extravasation of tumor cells. We next documented and quantified the extravasation process (that can be assessed at different stages from 3 to 24 hpi) by tracking the position of TCs with regards to the vasculature as previously described (Follain et al., 2018 ; Follain et al., 2021) (Figure 4A). When probing metastatic extravasation of mechanically-tuned CTCs (24 hpi) (Figures 4B-C), we found that siMYH9-treated tumor cells, in which elasticity was significantly reduced while viscosity remained unaltered, had the same extravasation potential as controls while all the other conditions in which viscosity was significantly reduced (siVIM, siCAV1, siLMNA) displayed reduced ratios of extravasated cells (Figures 4D, Fig.S4). While this strongly suggested that low-viscosity tumor cells had reduced extravasation efficiencies, we next used live imaging-based morphometric analysis at the early stage of extravasation (3 hpi) to better understand how cell viscosity could be involved in the process. As we previously described in the past with high-resolution imaging (Follain et al., 2018 ; Follain et al., 2021 ; Karreman et al., 2023), arrested tumor cells exploit the intraluminal remodeling activity of endothelial cells to perform extravasation. When probing mechanically-tuned CTCs, we realized that CTCs’ whose viscosity had been reduced (siLMNA, siVIM, siCAV1) were significantly less likely to engage such endothelial remodeling which can easily be detected and quantified (Figures 4E, S5). In contrast, CTCs with higher viscosities (siCTRL and siMYH9) were frequently enclosed and pocketed by endothelial projections (Figures 4F-G). This consequently increased the number of intravascular cells that remained unengaged with the endothelium for low viscosity tumor cells (Figure S6A). This difference wasn’t caused by differential cell distribution within the caudal plexus, as the exact same trend could also be seen when quantifying only the events that occurred in large vessels (Figure S6B). While pocketing events could be seen in all vascular subregions of the caudal plexus (Figures 4H, S5, S6C), we noticed that fully-extravasated cells could also frequently be seen around ISVs (Figure 4I) or around the complex architecture of the CV/CPCs (Figure S6D). As these vascular environments are the most mechanically-constraining ones (SVs), we wondered whether they would also offer optimal conditions for successful metastatic extravasation. Indeed, we found that a quarter of tumor cells completed extravasation within 3 hpi within these regions (ISVs, CV/CCPs), which is significantly more than in other non-constraining regions (DA, AVJs) where these events remained extremely rare (Figure 4J). Altogether, these results suggest that cell viscosity could impact metastatic extravasation on multiple levels, with low viscosity granting easier access to early extravasation vascular regions (SVs) but yet at the same time causing a significant and systematic delay in activation of endothelial remodeling-driven extravasation (Fig. 4K).

**Figure 4:**
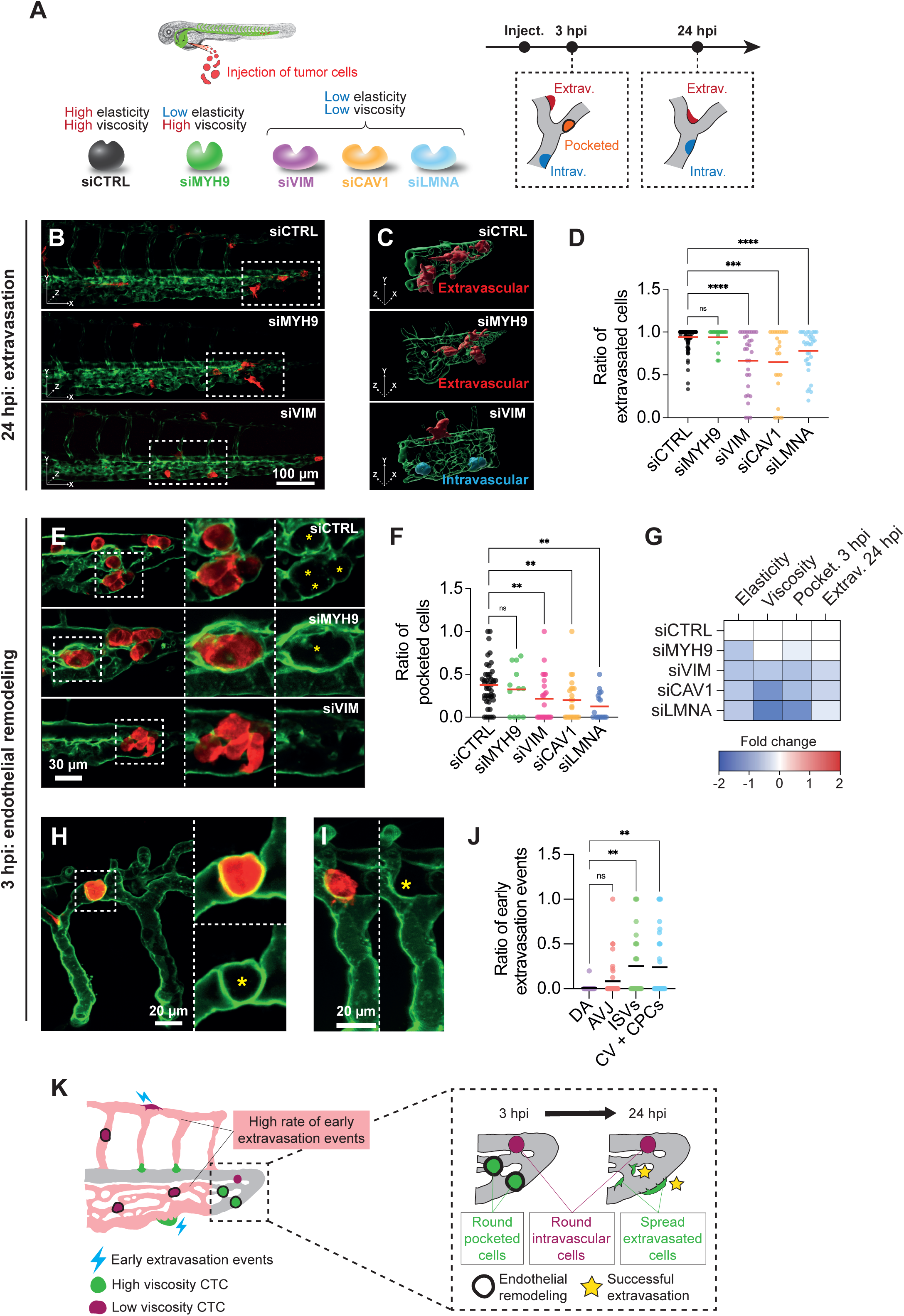
Tumor cell viscosity tunes metastatic extravasation. (A) Workflow for assessing extravasation abilities of mechanically-altered tumor cells in the ZF embryo model. (B) Z-projections of confocal stacks of siRNA-treated D2A1 tumor cells arrested in the caudal plexus of ZF embryo 24 hours post-injection. (C) 3D renders of framed areas in (B) and discrimination of intravascular and extravascular cells. (D) Quantification of ratio of extravasated cells 24 hours post-injection of mechanically-altered tumor cells. (N = 12 (siCTRL) ; 3 (siMYH9, siVIM, siCAV1, siLMNA), n (embryos) = 128 (siCTRL) ; 18 (siMYH9) ; 30 (siVIM) ; 23 (siCAV1) ; 33 (siLMNA)) (Kruskal-Wallis, p-values = 0.6184 (siCTRL vs siMYH9) ; <0.0001 (siCTRL vs siVIM) ; 0,0003 (siCTRL vs siCAV1) ; <0.0001 (siCTRL vs siLMNA)). (E) Z-projections of confocal stacks of siRNA-treated D2A1 tumor cells arrested in the caudal plexus of ZF embryo 3 hours post-injection. Yellow stars indicate tumor cell-containing endothelial pockets. (F) Quantification of ratio of pocketed cells 3 hours post-injection of mechanically-altered tumor cells. (N = 4 (siCTRL) ; 2 (siMYH9, siVIM, siCAV1, siLMNA), n (embryos) = 45 (siCTRL) ; 12 (siMYH9) ; 24 (siVIM) ; 23 (siCAV1) ; 17 (siLMNA)) (Kruskal-Wallis, p-values = 0.5202 (siCTRL vs siMYH9) ; 0.0080 (siCTRL vs siVIM) ; 0,0076 (siCTRL vs siCAV1) ; 0.00015 (siCTRL vs siLMNA)). (G) Fold changes in ratios of elasticity, viscosity, pocketed cells 3 hpi and extravasated cells 24 hpi for mechanically-altered tumor cells. (H) Z-projection of confocal stack of D2A1 tumor cells pocketed in the ISV vascular subregion of the caudal plexus of ZF embryo 3 hours post-injection. Yellow star indicates tumor cell-containing endothelial pocket. (I) Z-projection of confocal stack of D2A1 tumor cells extravasated in the ISV vascular subregion of the caudal plexus of ZF embryo 3 hours post-injection. Yellow star indicates extravascular location of the tumor cell. (J) Quantification of ratio of early extravasation events in different vascular subregions of the caudal plexus of the ZF embryo 3 hours post-injection of siCTRL-treated D2A1 tumor cells. (N = 4, n (embryos) = 26, 39, 27 and 34 for the DA, the AVJ, the ISVs and the CV+CPCs respectively) (Kruskal-Wallis, p-values = 0.1663 (DA vs AVJ) ; 0.0070 (DA vs ISVs) ; 0.0073 (DA vs CV+CPCs) ; 0.0586 (AVJ vs ISVs) ; 0.0881 (AVJ vs CV+CPCs). (K) Graphical summary of the impact of cell viscosity on tumor cell extravasation.

Having linked the viscosity of CTCs to endothelial remodeling-driven extravasation, whose occurrence is high (in ZF embryos or during mouse brain metastasis (Follain et al., 2018 ; Follain et al., 2021 ; Karreman et al., 2023)), we wondered whether this extravasation process (Figure S6E) was accompanied by cell deformation which accompanies classical diapedesis/transmigration extravasation events (Roberts et al., 2021). When comparing cell shapes of tumor cells at different stages of the extravasation process (3 hpi), we found no differences in volume between intravascular, pocketed and extravasated cells (Figure S6F). Interestingly, increased elongation (Figure S6G) and flatness (Figure S6H) of tumor cells was detected only in cells that had fully completed extravasation, likely due to post-extravasation cell spreading, but remained unchanged when comparing arrested intravascular cells and pocketed ones in the process of being extravasated. This morphometric analysis further suggests that volume or shape changes are not prerequisites for successful endothelium-mediated extravasation. Altogether, our results demonstrate for the first time that tumor cell viscosity impacts the extravasation step of metastasis. While early extravasation is more likely to occur in constraining vascular regions accessible to low-viscosity tumor cells, these cells are less efficient at triggering the endothelial remodeling-driven extravasation process (Figure 4K).

### The viscoelastic profile of CTCs has a lasting impact on post-extravasation metastatic outgrowth

Encouraged by the identification of viscosity as a key physical property of metastatic extravasation, we next sought to investigate its impact on seeding, extravasation and metastatic outgrowth in a syngeneic experimental metastasis mouse model (Figure 5A). First, early (1hpi) *ex vivo* lung imaging of i.v. injected tumor cells (tail vein) (Figure 5B) was combined with morphometric analysis of their reconstructed 3D shapes (Figure S7A) and revealed that mechanically-tuned tumor cells successfully reached constraining lung capillaries and displayed similar elongation and flatness factors (Figures S7B-C). Interestingly, microcirculation within the mouse lung vasculature appeared to be extremely constraining for tumor cells as they displayed significantly increased elongation and flatness factors compared to values that could be seen in the zebrafish embryo vasculature (Figures S7D-E). Interestingly, we found a 2-fold increase in tumor cell density within the mouse lung for low viscosity tumor cells (siVIM) compared to high viscosity cells (siCTRL, siMYH9) (Figure 5C) suggesting that, alike in ZF embryos, reduced viscosity grants access to constraining vessels and thus favors organ seeding. We then exploited this *ex vivo* model to track and quantify metastatic extravasation of i.v. injected mechanically-tuned tumor cells 24 hpi (Figure 5D). In line with the results obtained in the zebrafish embryo model, we found that extravasation of low viscosity tumor cells (siVIM) was significantly reduced (Figures 5E, S7F). Neither cell volume (Figure S7G), cell elongation (Figure S7H) nor cell flatness (Figure S7I) were perturbed when assessing intravascular and extravasated cells. We fully exploited this mouse model to track metastatic outgrowth by imaging and quantifying area of growing metastatic foci 10 dpi (Figure 5F). Confident that our mechanical tuning remained efficient 7 days post-transfection (Figure S7J-L), we observed that low viscosity tumor cells (siVIM) displayed increased outgrowth of established metastatic foci (Figure 5G). This suggests, for the first time, that the mechanical contribution of CTCs to the different steps of the metastasis cascade isn’t unilateral with regards to its success. Indeed, while high viscosity tumor cells have enhanced extravasation abilities, such property does not necessarily translate into increased metastatic outgrowth.

**Figure 5:**
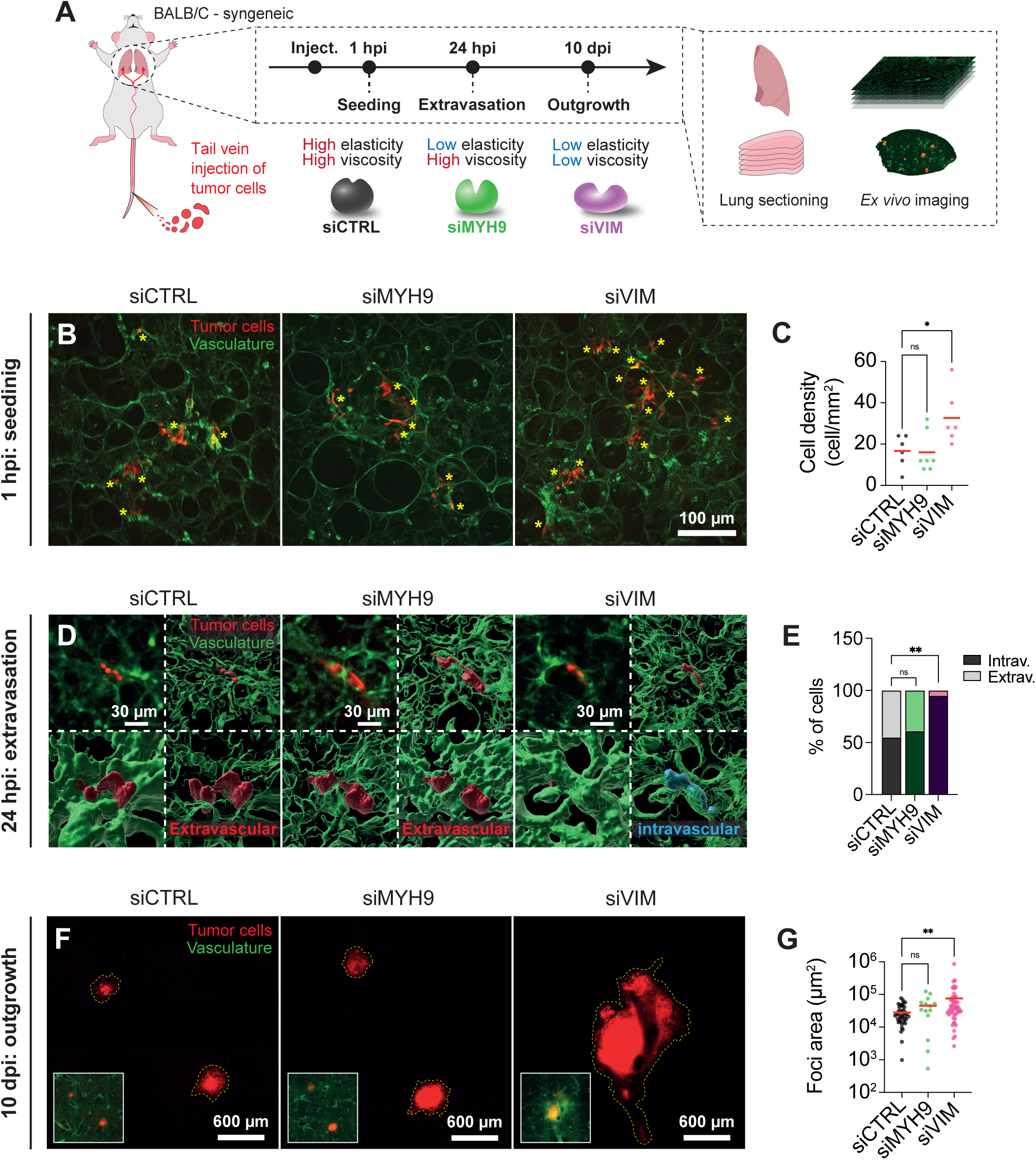
The viscoelastic profile of CTCs has a lasting impact on post-extravasation metastatic outgrowth. (A) Injection of mechanically-altered tumor cells in the mouse model (tail vein) for assessing seeding, extravasation and metastatic outgrowth in the mouse lungs with *ex vivo* imaging. (B) Z-projections of confocal stacks of mechanically-altered D2A1 tumor cells arrested in the mouse lung vasculature 1-hour post-injection. (C) Quantification of density of arrested mechanically-altered tumor cells found in the mouse lung vasculature 1-hour post-injection. (N = 1, n (fields of view) = 6 (siCTRL) ; 7 (siMYH9) ; 6 (siVIM)) (Kruskal-Wallis, p-values = 0.9084 (siCTRL vs siMYH9) ; 0.0361 (siCTRL vs siVIM)). (D) Z-projections of confocal stacks of mechanically-altered D2A1 tumor cells arrested in the mouse lung vasculature 24 hours post-injection and 3D renders for discrimination of intravascular and extravascular cells. (E) Quantification (in %) of intravascular and extravascular mechanically-altered D2A1 tumor cells in the mouse lung vasculature 24 hours post-injection. (N = 2, n (cells) = 22 (siCTRL) ; 18 (siMYH9) ; 19 (siVIM)) (Fischer’s exact test performed with raw number of cells, p-values = 0.7547 (siCTRL vs siMYH9) ; 0.0048 (siCTRL vs siVIM)). (F) Z-projections of tiled confocal stacks of metastatic foci originating from mechanically-altered tumor cells 10 days post-injection in the mouse lung vasculature. (G) Quantification of metastatic foci area originating from mechanically-altered tumor cells 10 days post-injection in the mouse lung vasculature (N = 4, 4 and 3 for siCTRL, siVIM and siMYH9 respectively, n = 41, 47 and 14 for siCTRL, siVIM and siMYH9 respectively) (Kruskal-Wallis, p-values = 0.0077 (siCTRL vs siVIM) ; 0.0983 (siCTRL vs siMYH9). (G) Graphical summary of the impact of viscoelasticity of tumor cells on post-extravasation metastatic outgrowth.

## Discussion

In this work, we demonstrate for the first time that cell viscosity is as a fundamental biomechanical property that affects hematogenous dissemination (circulation, arrest and extravasation) of circulating tumor cells, by enabling CTC entry and lodging in constraining capillary-sized vessels when it is low and favoring endothelial remodeling-assisted extravasation when it is high. Our results also hint at cell mechanics possibly having a lasting regulating impact on the post-extravasation fate of tumor cells, with low viscosity cells displaying increased metastatic outgrowth. Altogether, our results reinforce the hypothesis that tumor cells likely rely on their mechanical profiles to successfully complete the metastatic cascade (Gensbittel et al., 2021). On one hand, they exploit high-viscosity states to efficiently extravasate and colonize extravascular environments. On the other hand, they capitalize on low viscosity to reach and survive in constraining vascular regions, yet with reduced extravasation potential. Alternatively, one cannot exclude that tumor cells tune their mechanical profiles to adapt to the situation and ensure metastatic success.

In this study, we exploited polyacrylamide elastic beads as a tool to model CTCs and measure the stress they encountered *in vivo* when subjected to classical experimental metastasis schemes in the zebrafish embryo. Polyacrylamide elastic beads are capable of emulating the physical properties of both cells (Girardo et al., 2018) and tissues (Wagner et al. 2019) and thus bear enormous potential in mechanobiology. In this study, we exploited two major advantages of these beads. First, the beads were utilized as simplified models of CTCs with accurate and tunable mimicking of the diameter and elasticity of tumor cells. They exhibit near-elastic behavior, as they undergo almost instantaneous deformation with a relaxation time at the millisecond scale (Reichel et al., 2024) and as their mechanical properties remain time-independent across a broad range of time scales, spanning from milliseconds to seconds (Girardo et al., 2018). As such, they allowed us to highlight and reinforce to what extent cell mechanics and size control intravascular behavior. Indeed, by analyzing circulation patterns of various elastic bead populations, we identified reduced diameter and elasticity as properties that favor transit through occlusion sites and grant access to small constraining vessels.

Second, as force sensors, they enable the accurate evaluation of stresses at the cellular level via analysis of their deformation (Taubenberger et al., 2019 ; Träber et al., 2019 ; Zhang et al., 2023). By measuring the mechanical stress that these near-elastic objects encounter at various intravascular arrest sites and comparing their deformation to the ones of viscoelastic tumor cells, we pinpointed to the importance of viscosity in controlling CTC arrest. To our surprise, we found that the pressure sustained by beads was similar in all vascular subregions of the zebrafish embryo caudal plexus, irrespective of vessel diameter or whether the sites constituted hotspots of occlusion. This suggests that elastic beads are poorly constrained once arrested, very likely because vessels reshape in response to their presence, rather than the elastic beads getting deformed by the constraining vessels and associated hemodynamics (Figure 2B-C). Intravascular endothelial deformation occurs rapidly around arrested objects (Lam et al., 2010) and even mediate metastatic extravasation (Follain et al., 2018 ; Follain et al., 2021 ; Karreman et al., 2023). Assessing to what extent endothelial deformation, through its own mechanical profiles that one can probe using biosensor imaging (Lagendijk et al., 2017), is likely to inform onendothelial cell mechanics, in addition to tumor cells ones, in mediating efficient intravascular arrest. The elasticity level of tumor cells (0.8 kPa) is, in theory, sufficiently high for them to resist mechanical pressure within the intravascular environment and induce deformation of vessels around them instead of undergoing deformation themselves, as implied by the behavior of our cell-mimicking beads. Yet, our 3D shape factor analysis revealed that arrested tumor cells displayed significantly more deformation compared to arrested elastic beads within the intravascular environment. Two parameters could explain this observation: 1) Tumor cells could at some point actively adapt their shape while engaging into migratory behavior and in doing so, release the pressure that their presence would exert on the vasculature. 2) Their viscosity could rapidly take over after elastic behavior and cause them to passively deform to extensive levels when faced with a constraining intravascular environment.

When dynamically probing the deformation of elastic beads and viscoelastic tumor cells, we observed that our polyacrylamide elastic beads, which are characterized by a cross-linked network, remain occluded until a critical pressure is reached and undergo rapid (<1sec) elastic deformation to enter constraining vessels. On the contrary, tumor cells display a progressive and slow viscous-like deformation as they enter, creep and move within constraining vessels. These observations suggest, for the first time, that cellular viscosity is a key determinant of CTCs’ ability to enter constraining vessels and highlight the need to design, in the future, tunable viscoelastic artificial objects. Controlling the viscous component of the beads, in addition to the existing control of elasticity, will allow greater flexibility in experimental design to study the impact of cell mechanics on both short and long-term deformation dynamics within confining vessels. To this end, viscoelastic beads with elasticity similar to that of CTCs and a significant viscous component could be generated using hydrogels that form a physically entangled network (Oyen, 2014). Such tools are likely to become available in the near-future and will provide invaluable insights as near-perfect force-sensing tools.

Aiming at further dissecting the relative contribution of elasticity over viscosity of tumor cells in controlling their intravascular fate, we targeted several components likely to affect one or the other or both. While knocking-down lamin A/C, vimentin or caveolin-1 led to combined decreases in both elasticity and viscosity, depletion of myosin IIA led to decreased elasticity only. These effects were all in line with the known involvements of these proteins in cell mechanics (Vahabikashi et al., 2022 ; Janmey et al., 1991 ; Lolo et al., 2022 ; Urbanska et al., 2023 ; Chugh and Paluch, 2018) and constituted yet another demonstration that the cytoskeleton represents a key regulator of the viscoelastic profile of cells (Urbanska and Guck, 2024). This begs the question of how tumor cells may exploit their cytoskeleton to adapt their mechanical properties. Of note, the intermediate filament protein vimentin, which we found to be strongly linked to cell viscosity throughout our study, is notoriously regulated differently across the epithelial-to-mesenchymal transition (EMT) spectrum, implying that EMT programs could be intricately linked to mechanical phenotype changes in tumor cells (Lorenz et al., 2023), as discussed later.

In line with the identification of viscosity as a major enabler of CTC escape from occlusion sites, we observed that viscosity was the only mechanical parameter that significantly correlated with successful arrest in small constraining vessels. This was not the case for elasticity, reinforcing the need to dissect which cellular components control one (viscosity) or the other (elasticity). Interestingly, recent evidence suggests that cytoplasmic viscosity bears potential in as a mechanical marker for metastatic tumor cells (Dessard et al., 2024). Yet, one cannot fully exclude a lesser contribution of elasticity in the process of CTCs seeding capillary-sized vessels, as we also found low elasticity to facilitate elastic bead entry in small vessels.

When further dissecting the contribution of cellular viscosity to the intravascular arrest of CTCs, we realized that CTCs whose mechanical properties allowed them to reach small-sized vessels could potentially benefit from undergoing rather quick extravasation. Indeed, accurate assessment of such dynamics that is offered by the zebrafish embryo model showed that a quarter of arrested tumor cells were already extravasated 3 hours after injection. Probing the behavior of such vessels is highly relevant as they closely mimic capillary beds in metastatic organs where CTCs have often been found to arrest (Kienast et al., 2010 ; Entenberg et al., 2015 ; Headley et al., 2016). This further suggests that low viscosity CTCs evolved means to quickly extravasate from constraining and hostile vessels, for two reasons: on one hand, blood flow interruption that is triggered by an arrested CTC in a capillary is often counteracted by the need to rescue blood perfusion, which vessels easily do through endothelial remodeling (Lam et al., 2010); on the other hand, performing extravasation quickly would be beneficial to tumor cells as the intravascular environment and its hemodynamic forces are hostile for them (Follain et al., 2020 ; Moose et al., 2020). Yet, although constraining vessels seem prone to fast extravasation events, they constitute an obstacle to high viscosity tumor cells. In addition, our data strongly suggests that low viscosity CTC are less likely to trigger extravasation-prone endothelial remodeling. This raises the question as to which viscosity-related mechanism (and not elasticity) is likely to stimulate intravascular remodeling. While we previously found this process to be mechanosensitive to hemodynamic forces (Follain et al., 2018 ; Follain et al., 2021) and to the secretion of MMP9 (Karreman et al., 2023), one could hypothesize that stiffer (more viscous) objects are more likely to transmit forces to the surrounding endothelial cells and, doing so, force the much-needed remodeling. In the end, it seems likely that some tumor cells are capable of hijacking a mechanism meant to recanalize occluded microvessels (Lam et al., 2010 ; Karreman et al., 2023), and that the most mechanically inconvenient tumor cells (bigger, stiffer, more viscous) could be more likely to trigger this mechanism. Yet, cell-mimicking polyacrylamide elastic beads do not trigger endothelial remodeling and do not undergo extravasation. Such phenomenon requires molecular adhesion machineries that favor interactions between membranes of endothelial and tumor cells and activation of ligand-receptor crosstalk (Gassmann et al., 2009 ; Osmani et al., 2019). It is tempting to speculate that the inner mechanics of cells makes them more likely to activate such receptors as it has been shown for immune cells during the immunological synapse formation for example (Liu et al., 2021 ; Tello-Lafoz et al., 2021 ; Lei et al., 2021).

While low viscosity profiles of tumor cells impair the extravasation potential, as seen both in the zebrafish embryo and in experimental lung metastasis in a syngeneic mouse model, the latter allowed us to show that low viscosity tumor cells gave rise to faster-growing metastatic foci. Vimentin was recently also shown to repress experimental lung metastasis model in the mouse model (Grasset et al., 2022), in line with our results, raising the question of linking EMT (epithelial-mesenchymal transition) states of metastatic cells with their mechanical profiles. While recent evidence suggests that circulating tumor cells that successfully reach distant organs display a wide spectrum of epithelial, mesenchymal and hybrid phenotypes, it is tempting to speculate that 1) similar behavior would apply to their mechanical profiles, where CTCs would display very heterogenous elasticity and viscosity profiles and that the latter 2) are tightly linked to such EMT state. Interestingly, we found that arrested tumor cells appeared to be much more elongated and flattened in the mouse lung vasculature compared to tumor cells arrested in large and even small vessels of the zebrafish embryo shortly after injection (1 hour). Thus, we can reasonably speculate that cell viscosity, which we found to be a critical determinant of large vs small vessel entry/arrest in the zebrafish embryo, also strongly impacts the ability of CTCs to reach the capillaries that constitute the mouse lung vasculature. Low viscosity (siVIM) CTCs could circulate and arrest in greater numbers in this constraining architecture, which could compensate for the extravasation defect that we found for these cells and lead to the increased outgrowth phenotype that we and others (Grasset et al., 2022) have described.

With obvious links to their EMT profile, it is likely that the mechanical phenotype of tumor cells is linked to their fitness to proliferate after extravasation and thereby colonize a metastatic organ, with soft tumor cells possibly displaying increased stemness and tumorigenicity compared to stiffer ones (Lv et al., 2020). It is tempting to suggest that probing the molecular (EMT status) and the mechanical profiles (elasticity and viscosity) of patient-derived tumor cells, especially CTCs, is likely to provide useful prognostic markers. Interestingly and in line with our observations, recent evidence suggest that viscosity might be a strong indicator of the metastatic potential of tumor cells (Dessard et al., 2024). This once again begs the question of whether tumor cells might also dynamically adjust their mechanical properties based on their environment in order to progress along their metastatic journey. In a similar way to clustering (Aceto et al., 2014), hybrid EMT phenotypes which might allow tumor cells to perform cytoskeletal switches and adjust their viscoelastic profile (Lorenz et al., 2023) were found to grant enhanced metastatic potential to tumor cells (Yu et al., 2013; Padmanaban et al., 2019). Further investigation is required to better dissect and understand whether cell mechanics truly impacts the post-extravasation fate (dormancy vs proliferation) of tumor cells and whether mechanical switches truly occur in tumor cells as they travel between intravascular and extravascular environments.

Overall, our work highlights viscosity as a fundamental property of circulating tumor cells with strong impact on circulation, arrest, extravasation and potentially even their fitness to perform metastatic outgrowth. We found low viscosity to be beneficial for microcirculation and lodging in capillary-sized vessels but detrimental for endothelium-assisted extravasation. All-in-all, our results reinforce the notion that successful metastasis is achieved by tumor cells capable of optimally exploiting their mechano-molecular profile or their ability to dynamically adjust it along the way.

## Materials and methods

### Cell Lines / Cell culture

D2A1 Cells (CVCL_0I90). Mouse Mammary Carcinoma (BALB/c female).

Culture conditions: 37°/5% CO_2_. DMEM HG with 5% NBCS, 5% FBS, 1% NEAA-MEM, 1% Penstrep. Authentication: Injection in the nipple of mammary gland of BALB/c mice lead to mammary tumor. Cells do not show contamination to mycoplasma.

JIMT-1 BR3 Cells (CVCL_2077). Human Ductal Breast Carcinoma (Female) Highly Metastatic in the Brain.

Culture condition: 37°/5%CO_2_. DMEM HG with 10% FBS, 1% Penstrep. Authentication: Intracardiac injection in nude mice (NU/NU) lead to cerebral metastasis. Cells do not show contamination to mycoplasma.

1675 Cells (WM115, CVCL_0040). Human Melanoma (Female).

Culture condition: 37°/5%CO_2_. DMEM HG with 10% FBS, 1% Penstrep. Cells do not show contamination to mycoplasma.

A431 Cells (CRL_1555). Human Skin Epidermoid Carcinoma (Female).

Culture conditions: 37°/5% CO_2_. DMEM HG with 10% FBS, 1% Penstrep. Cells do not show contamination to mycoplasma.

4T1 Cells (CVCL_0125). Mouse Mammary Gland Carcinoma (BALB/c Female).

Culture condition: 37°/5%CO_2_. RPMI 1640 with 10% FBS, 1% Penstrep. Authentication: Injection in the nipple of mammary gland of BALB/c mice lead to mammary tumor. Cells do not show contamination to mycoplasma.

ZMEL1 Cells (Zebrafish melanoma).

Culture condition: 28°/5% CO_2_. DMEM HG, 10% FBS, 1% Penstrep. Cells do not show contamination to mycoplasma.

### Cell line engineering

For in vivo imaging in zebrafish embryos, D2A1 cells were engineered to express a Lifeactin-GFP or a Lifeactin-tdTomato fusion protein. Briefly, the Lifeactin-GFP or Lifeactin-tdTomato DNA fragment was inserted in a pLSFFV-Ires-Puromycin lentiviral vector to generate pLSFFV-LifeActin-GFP-Ires-Puro and pLSFFV-LifeActin-tdTomato-Ires-Puro lentiviral vectors used to transduce D2A1 cells. Vimentin-overexpressing D2A1 cells were obtained similarly with a mouse vimentin DNA fragment kindly provided by Sandrine Etienne-Manneville. Lentivirus particles production and transduction procedures were performed as described previously (Hyenne et al., 2019).

### siRNA transfection

D2A1 stably expressing LifeAct-tdT or LifeAct-GFP were grown as previously described (Shibue et al., 2013), in DMEM with 4.5 g/l glucose (Dutscher) supplemented with 5% FBS, 5% NBCS, 1% NEAA and 1% penicillin-streptomycin (Gibco). siRNAs were transfected into D2A1 cells using Lipofectamine RNAiMAX (Invitrogen) following the manufacturer’s instructions. Experiments were performed 72 h post-transfection.

The following siRNA sequences were used:

siCTRL: Dharmacon ON-TARGET plus Non-targeting Pool (D-001810-10-20) or Sigma-Aldrich custom-made non-targeting sequences: GCAAAUUAUCCGUAAAUCA [dt][dt] and UGAUUUACGGAUAAUUUGC [dt][dt]

siLMNA: Dharmacon ON-TARGETplus Mouse Lmna (16905) siRNA – SMARTpool (L-040758-00-0010) or Sigma-Aldrich custom-made non-targeting sequences: UGACCAUGGUUGAGGACAA [dt][dt] and UUGUCCUCAACCAUGGUCA [dt][dt]

siVIM: Dharmacon ON-TARGETplus Mouse Vim (22352) siRNA – SMARTpool (L-061596-01-0010) or Sigma-Aldrich custom-made non-targeting sequences: AGGCCAAGCAGGAGUCAAA [dt][dt] and UUUGACUCCUGCUUGGCCU [dt][dt]

siMYH9: Sigma-Aldrich custom-made sequences GACAAAGGUUCGAGAGAAA[dt][dt] and UUUCUCUCGAACCUUUGUC[dt][dt]

<u>siCAV1:</u> Dharmacon ON-TARGETplus Mouse Cav1 (12389) siRNA – SMARTpool (L-058415-00-0010)

### Micropipette aspiration assay

Cells were trypsinized and kept in PBS during MPA experiments. Measurements were performed on a custom-made MPA setup equipped with a camera and connected to a computer. Capillaries used for the aspirations were micro-forged to set their opening diameter to 3,5 µm. A vertical water column displacement of 10,5 cm was used to induce suction of the cells in the capillary. Movies of cell aspirations were recorded at 5 frames per second and edited in FIJI software (Schindelin et al., 2012). Movies were cropped, rotated and edited to have the entering of the cell in the capillary start precisely at the first frame of the recordings and progress vertically at an angle of 90°.

The “orthogonal view” tool was used to generate the curve representing the cell entering the capillary in function of time. The segmented line tool was used to manually trace the curves and generate the associated coordinates. Obtained coordinates were opened in IGOR Pro7 (WaveMetrics software). Coordinates values were converted from pixels to microns for the Y wave (entering of the cell in the capillary) and from pixels to seconds for the X wave (time elapsed). Minor adjustments to the coordinates were done to make all curves start at the origin of the graph (the coordinates of the first point of the curves were set at (0;0). Finally, curve-fitting was performed to extract elasticity and viscosity coefficients of aspirated cells using a rheological model for micropipette aspiration of biological viscoelastic droplets described previously (Guevorkian et al., 2010).

### Real-Time Deformability Cytometry

Real-time deformability cytometry (RT-DC) was performed as described elsewhere (Otto et al., 2015 ; Herbig et al., 2017 ; Rosendahl et al., 2018) using an AcCellerator (Zellmechanik, Dresden, Germany) at 37 °C. In brief, D2A1 cells were detached from cell culture plates using trypsin and kept on a roller in the culture medium in an incubator (37°C) for 30 min to reduce their surface/volume ratio. Subsequently, the cells were strained through a 40 µm strainer, centrifuged (200xg, 5 min), and resuspended in PBS containing 0.5% (w/v) methylcellulose (measurement buffer). RT-DC was performed using microfluidic chips with channels of 30 × 30 μm cross-section. The total flow-rate was set to 0.240 μL/s, and acquisition was performed using ShapeIn software (Zellmechanik, Dresden, Germany). The Young’s modulus was calculated from deformation and cell area using ShapeOut software (Müller, E. O’Connell, M. Schlögel, Shape-Out version 2.9.3: Graphical user interface for analysis and visualization of RT-DC data sets, GitHub (2022) (https://github.com/ZELLMECHANIK-DRESDEN/ShapeOut2) after filtering with the following setting: cell area parallel to flow 80–1000 μm^2^ (removal of cell debris/multicellular clusters), porosity 1.00–1.05 (removal of damaged cells).

### Zebrafish handling

*Tg(fli1a:eGFP)* and *Tg(kdrl:Hsa.HRAS-mCherry)* Zebrafish (*Danio rerio*) were crossed to obtain the embryos used in the experiments. Embryos were maintained in Danieau 0.3X medium (17,4 mM NaCl, 0,2 mM KCl, 0,1 mM MgSO_4_, 0,2 mM Ca(NO_3_)_2_) buffered with HEPES 0,15 mM (pH = 7.6), supplemented with 200 μM of 1-Phenyl-2-thiourea (Sigma-Aldrich) to inhibit the melanogenesis, as previously described (Goetz et al., 2014).

### Polyacrylamide bead production

Alexa Fluor™ 488-labeled polyacrylamide elastic beads were produced and analyzed as previously described (Girardo et al., 2018). Briefly, a flow-focusing PDMS-based microfluidic chip was used to produce polyacrylamide pre-gel droplets in fluorinated oil (3M™ Novec™ 7500, Iolitec Ionic Liquids Technologies GmbH) containing ammonium Krytox^®^ surfactant (2.4%w/v) as an emulsion stabilizer, 0.4%v/v N, N, N′, N′ - tetramethylethylenediamine (TEMED) (Sigma-Aldrich) as a catalyst and 0.1%w/v acrylic acid N-hydroxysuccinimide ester (NHS) (Sigma-Aldrich) as a compound to include NHS functional groups into the final gel meshwork for binding of Alexa Fluor™ 488 hydrazide. The pre-gel mixture contained acrylamide (40%w/v) (Sigma-Aldrich) as a monomer, bis-acrylamide (2%w/v) (Sigma-Aldrich) as a crosslinker, ammonium persulphate (0.05%w/v) (Sigma-Aldrich) as a radical initiator and Alexa Fluor™ 488 hydrazide (2 mg/ml) (Thermo Fisher) diluted in 10mM Tris buffer (pH = 7.48). Total monomer concentration in the pre-gel mixture together with the droplet diameter was fine-tuned to obtain beads with well-defined diameter and elasticity. After in-drop polymerization, the beads were washed and resuspended in 1xPBS (pH = 7.4). Elasticity was measured by Atomic Force Microscopy (AFM) indentation and Real Time-Deformability Cytometry (RT-DC). Bead polydispersity in elasticity was reduced by sorting them with fluorescence-activated cell sorting (FACS) resulting also in a reduction in diameter distribution. Here, the list of the beads used in the study reporting both mean diameter ± standard deviation and Young’s modulus (elasticity) ± standard deviation:

“Large beads”: diameter = 18.9 ± 0.4 µm, elasticity = 1.2 ± 0.4 kPa

“Small beads”: diameter = 10.9 ± 0.3 µm, elasticity = 0.9 ± 0.2 kPa

“Stiff beads”: diameter = 14.4 ± 0.4 µm, elasticity = 2.0 ± 0.2 kPa

“Soft beads”: diameter = 14.5 ± 0.6 µm, elasticity = 0.3 ± 0.1 kPa

“D2A1-mimicking beads”: 3 batches were used:

- diameter = 14.6 ± 0.5 µm, elasticity = 0.8 ± 0.1 kPa

- diameter = 14.3 ± 0.4 µm, elasticity = 0.8 ± 0.2 kPa

- diameter = 14.2 ± 0.4 µm, elasticity = 0.8 ± 0.1 kPa

Beads were stored in PBS at 4°C.

### Intravascular injections of beads and tumor cells in the zebrafish embryo

48-hour post-fertilization (hpf) *Tg(Fli1a:eGFP)* of *Tg(Kdrl:Hsa.HRAS-mCherry)* embryos were mounted in 0.8% low melting point agarose pad containing 650 μM of tricain (ethyl-3-aminobenzoate-methanesulfonate) to immobilize the embryos. D2A1 LifeAct-RFP or LifeAct-GFP cells were injected with a Nanoject microinjector 2 (Drummond) and micro-forged glass capillaries (25 to 30 μm inner diameter) filled with mineral oil (Sigma). Cell suspensions were prepared at approximately 40.10^6^ cells per ml for loading into the injection capillaries. Injection volume was initially set at 18nL but could be adjusted throughout injection sessions anywhere between 9nL and 27nL. Injections were targeted at the duct of Cuvier of the embryos and were performed under a M205 FA stereomicroscope (Leica), as previously described (Stoletov et al., 2010). The same protocol was used when injecting 1675, NIH3T3, A431, 4T1 and ZMEL1 cells. Similarly, injections of fluorescent polyacrylamide elastic beads were performed with similar concentrations of beads.

### Live recording of early circulation events in the zebrafish embryo

The earliest bead/cell circulation and arrest events following injection in the zebrafish embryo were captured with the M205 FA stereomicroscope (Leica) under which the injections were performed. The stereomicroscope was connected to a laptop equipped with Leica’s LAS AF software, allowing image/video acquisition. Exposure time was set to 200 ms in the appropriate channel for capturing the fluorescent signal of the beads or tumor cells, which resulted in a rate of acquisition of one frame every 0.43 s. Recordings of 5 minutes started right after embryo injection was performed. Once recordings of bead/cell circulation were complete, a single image was acquired in the appropriate channel for capturing the fluorescent signal of the embryo’s endothelium.

### Confocal imaging of injected zebrafish embryos

Confocal imaging was alternatively performed with an upright Leica TCS SP5 (20X objective, 0.75 NA) or an inverted Olympus Spinning Disk (30X objective, 0.8 NA). The caudal plexus region (around 50μm width) was imaged with a z-step of 2 μm in experiments aiming at locating and/or assessing the extravasation status of arrested tumor cells. The z-step was set to 0.25 µm in experiments aiming at performing 3D reconstruction and shape/volume analysis of arrested beads or cells.

### Ex vivo imaging of tumor cells in the mouse lung vasculature

BALB/C mice were injected in the tail-vein with siRNA-treated D2A1 tumor cells. 150.000 cells were injected for visualizing arrested cells in the lungs 1-hour post-injection, 500.000 cells were injected for the purpose of imaging extravasation events 24 hours post-injection, and 700.000 cells were injected for imaging growing metastatic foci 10 days post-injection. On the day of imaging, mice were injected with iFluor488-Wheat Germ Agglutinin (10µg, ref : AAT-25530) 5 minutes prior to sacrifice in order to stain the lung vasculature. During dissection, the thoracic cage was removed until the lungs were fully visible and muscles surrounding the trachea were also removed. A suture thread node was pre-made around the trachea but was left open prior to lung inflation. Using a 10 mL syringe with an 18-gauge needle inserted in the trachea, a 40°C mixture of low-melting agarose in PBS (4%) and medium at 1:1 volume was slowly injected until lungs were fully inflated. When lungs appeared inflated, the suture thread node was tightened to avoid medium leak and ice-cold PBS was poured on the lungs to solidify the agarose. Lungs were then dissected and placed in a tube containing ice cold PBS for 20 minutes. The different lobes were then cut using a vibratome in order to obtain 250 µm-thick slices for imaging.

Confocal imaging was performed using an inverted Olympus Spinning Disk. A 30X objective (0.8 NA) was used for acquiring stacks of arrested tumor cells shortly (1-2 hpi) after injection with a z-step of either 0.25 or 0.5 µm. A 10X objective (0.4 NA) was used for acquiring stacks of arrested tumor cells 24 hours after injection with a z-step of either 0.25 or 0.5 µm. A 10X objective (0.4 NA) was used with a scanning function to acquire large tiles of confocal stacks of entire lung sections with a z-step of 15 µm in order to find growing metastatic foci 10 days post-injection.

### Image processing and analysis

All image analysis of tumor cells in the zebrafish embryo model were performed in FIJI software (Schindelin et al., 2012).

<u>For early circulation and arrest of beads and CTCs</u>, the stereomicroscope still images of the embryos’ endothelium were duplicated and stacked to match the frame numbers of their corresponding video recordings of bead / CTC signal. Both channels were merged to visualize bead / CTC circulation and arrest in the blood vessels. Arrest events were manually counted and reported on a stereotyped reference map of the caudal plexus for heatmapping (See “heatmapping” section).

<u>For longitudinal tracking of shape changes of beads / CTCs at ISV entrances</u>, events of bead / CTC arrival at ISV entrance followed by successful entry were cropped and isolated from merged stereomicroscope recordings of bead / CTC circulation and arrest. Thresholding was used to generate binary mask stacks of bead / CTC shape in which aspect ratios were measured.

<u>For quantification of intravascular, pocketed and extravasated cells at 3- or 24-hours post-injection time points</u>, confocal z-stacks were carefully scrolled through to evaluate the extravasation status of arrested cells. Events were manually counted and reported on a stereotyped reference map of the caudal plexus for heatmapping.

<u>For analysis of bead / cell shape factors</u>, z-steps of confocal stacks were first adjusted to account for spherical aberrations using a plugin previously described (Diel et al., 2020), allowing faithful 3D reconstruction of beads / cells. “Gaussian blur”, “thresholding”, “make binary”, and “fill holes” functions were used to generate clean binary stacks of whole shape of beads / cells. 3D ImageJ Suite plugin (https://mcib3d.frama.io/3d-suite-imagej/) was then used to measure 3D ellipsoid shape descriptors of processed stacks.

Image analysis of tumor cells arrested in the mouse lung vasculature were performed in IMARIS software.

For analysis of shape factors of tumor cells arrested in the mouse lung vasculature (1-2 hpi), segmentations of cell shapes were performed, and elongation and flatness values were calculated directly from the built-in IMARIS shape descriptor tool that provides axis lengths of ellipsoid fits of the cells.

For assessing extravasation status of tumor cells arrested in the mouse lung vasculature (24 hpi), segmentation of both cells and vessels were performed and transparency of the vasculature was tempered with in order to discriminate intravascular and extravasated cells.

Image analysis of growing metastatic foci was performed in FIJI software. Brightness/contrast adjustments of z-projections of tiles of confocal stacks of entire lung sections were performed for optimal visualization of metastatic foci within the lung sections. Foci were detoured using the freehand selection tool and area of metastatic foci were measured using FIJI’s “measure” function.

### Bead deformation conversion to mechanical stress

Computational analysis-based cell-scale stress sensing (COMPAX) was performed as described previously (Träber et al., 2019). Beads used for this purpose went through extensive sorting steps in order to reduce the standard deviation of bead diameter as much as possible (+/– 0.4 µm) within the bead population, allowing for more accurate computation based on more accurate assumed non-deformed bead configurations. Simulations were also run multiple times for each bead, with a diameter distribution of assumed non-deformed beads reflecting the actual diameter distribution and standard deviation within the bead population injected and imaged.

### Heat-mapping

The heat-maps were generated using FIJI (Schindelin et al., 2012) and MATLAB (MathWorks) software as previously described (Follain et al., 2018). Events of interest (arrested / intravascular / pocketed / extravasated cells) were identified in embryos after careful analysis of the stereomicroscope recordings or confocal z-stacks with FIJI. The positions of these events were manually reported on a reference map of the stereotyped 2.5 days post-fertilization zebrafish embryo caudal plexus vasculature. Then, all the support images representing each embryo for one condition were put together in an image stack using ImageJ. The stack was read layer by layer in MATLAB and the dots representing event locations were automatically detected with the Hough circles function (Yuan-Liang Tang, Department of Information Management, Chaoyang University of Technology, Taichung, Taiwan) (https://fr.mathworks.com/matlabcentral/fileexchange/22543-detects-multiple-disks--coins--in-an-image-using-hough-transform) using the Circular Hough Transform based algorithm, giving in output the coordinates of the detected dots. Gaussian spots were then created at these coordinates. The amplitude of each Gaussian spot was equal to 1. The different layers of one condition were added to each other through a sum projection. The gaussian spot amplitudes of all layers were summed to produce the heatmap. The areas of the sum projection where the gaussian spot amplitudes are higher corresponds to high density areas of events. To produce the final heatmap, a custom colormap, inspired by the jet colormap, was applied to the sum projection. The colormap was then added on top of color-coded versions of the reference map initially used to report event locations in order to highlight densities of events in different vascular subregions of the zebrafish embryo caudal plexus.

### Western Blot

Protein extractions were performed with homemade RIPA buffer. Protein concentrations of the extracts were first quantified using a Bradford assay, and equal protein quantities were then loaded on 4-20% polyacrylamide gels (Biorad) or 4-20% Tris-Glycine gels (Introvigen) and run under denaturing conditions.

The following primary antibodies were used:

Anti-vimentin (D21H3), Cell Signaling

Anti-lamin A/C (sc-7292 and sc-20681), Santa Cruz

Anti-non-muscle myosin IIA (M8064), Sigma

Anti-caveolin-1 (D46G3), Cell signaling

## Acknowledgements

We thank all members of JGG’s team for their constant helpful discussions throughout this investigation. JGG is the coordinator of the NANOTUMOR Consortium, a program from ITMO Cancer of AVIESAN (Alliance Nationale pour les Sciences de la Vie et de la Santé, National Alliance for Life Sciences & Health) within the framework of the Cancer Plan (France). Work and people in the lab of JGG are mostly supported by the INCa (Institut National Du Cancer, French National Cancer Institute), charities (La Ligue contre le Cancer, ARC (Association pour la Recherche contre le Cancer), FRM (Fondation pour la Recherche Médicale)), the National Plan Cancer initiative, the Region Grand Est, INSERM and the University of Strasbourg. This work has been directly funded by the support of the Ligue Contre le Cancer (labelisation) and the association Ruban Rose. VG has been funded by INSERM, region Grand Est and La Ligue Contre le Cancer. GF was supported by La Ligue Contre le Cancer. LB is supported by FRM (Fondation pour la Recherche Médicale) (ECO202206015567). KU is funded by the Deutsche Forschungsgemeinschaft (DFG), project ID 467937258. VM is funded by a Ph.D. fellowship from the French Ministry of Science (MESRI) and by the Foundation ARC (Association pour la recherche contre le Cancer). JG acknowledges core funding by the Max Planck Society. The production and characterization of the polyacrylamide beads used in this study was supported by the European Union Horizon 2020 research and innovation programs No. 953121 (project FLAMIN-GO). RG is supported by FLAMIN-GO project. We thank Parth Patel, part of the TDSU Lab-on-a-chip systems at MPL, for the production of the master template and microfluidic chips used for the bead production. We thank Uwe Appelt and Markus Mroz of the Core Unit Cell Sorting and Immunomonitoring of the Nikolaus Fiebiger Centre for Molecular Medicine (NFZ) at the Friedrich Alexander University Erlangen-Nuremberg for their technical support in the bead sorting.

## Author contributions

Conceptualization: JGG, SH, NO, VG and JG. Methodology: OL, AL, SH, KU, DB, RG and SG. Investigation: VG, GF, KU, LB, OL, AF, VH, VM, MK and AD. Resources: RG and SG. Formal Analysis: VG and OL. Writing - Original Draft: VG, JGG, NO, GF, SH, MK, RG, SG and JG. Writing – Review & Editing: VG, JGG and NO. Supervision: JGG, JG, DB and VH. Funding Acquisition: JGG, NO, SG, DB and JG.

## Figure legends

**Figure S1 related to Figure 1:**
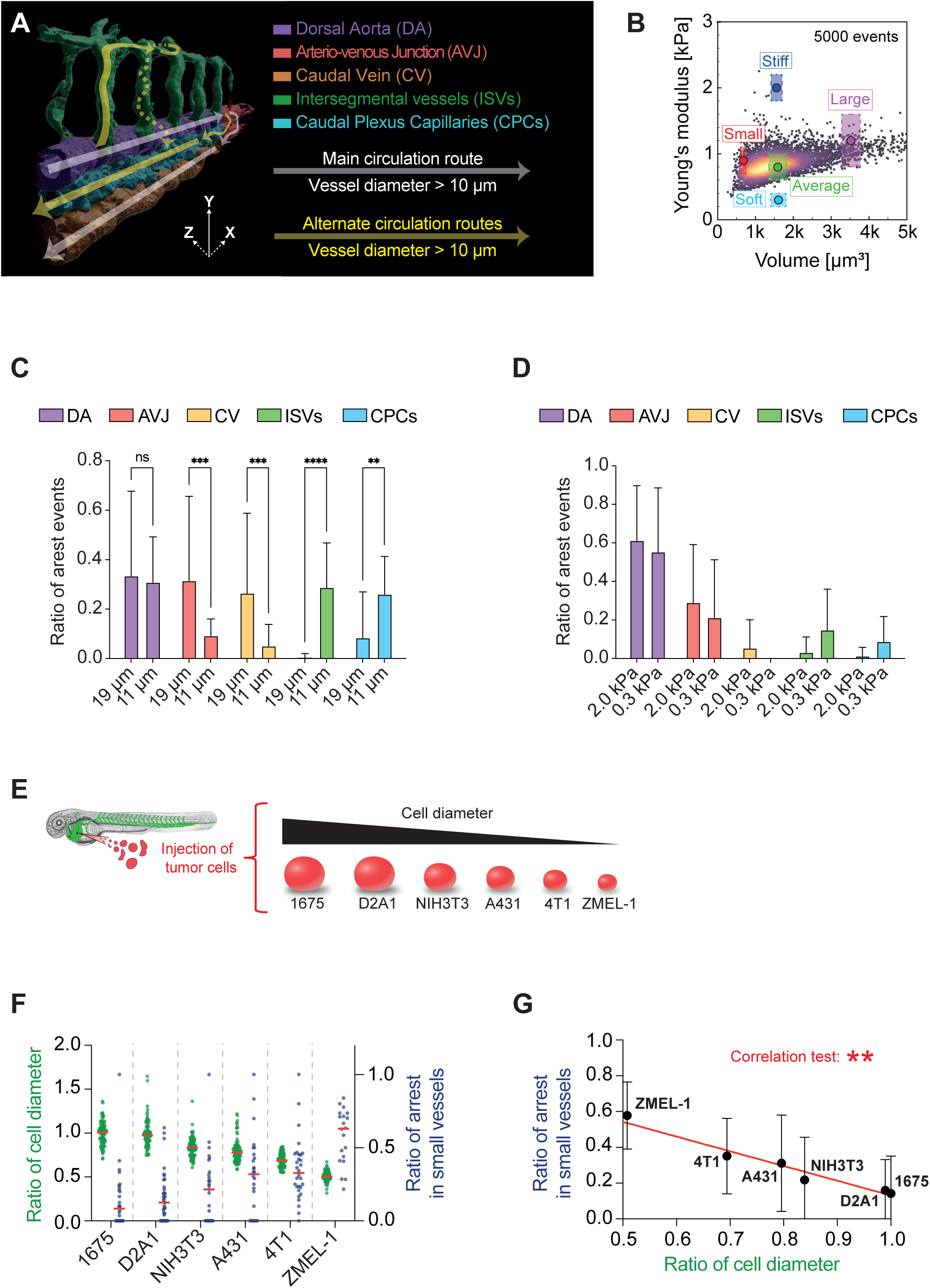
Small size and low elasticity of circulating objects facilitate lodging in constraining blood vessels. (A) 3D rendering of the zebrafish embryo caudal plexus. (B) Real Time-Deformability Cytometry (RT-DC) plot of D2A1 tumor cells and relative properties (size, elasticity) of polyacrylamide bead populations designed to model circulating tumor cells (N = 1, n (cells) = 5000). (C) Quantification of ratio of arrest occurring in each vascular subregions of the ZF embryo caudal plexus for large (19 µm) and small (11 µm) bead populations. (N = 2, n = 44 and 32 for large and small beads respectively) (2-way ANOVA, p-values = 0.9924 (DA) ; 0.0002 (AVJ) ; 0.0003 (CV) ; < 0.0001 (ISVs) ; 0.0048 (CPCs). (D) Quantification of ratio of arrest occurring in each vascular subregions of the ZF embryo caudal plexus for stiff (2.0 kPa) and soft (0.3 kPa) bead populations. (N = 2, n = 22 and 26 for large and small beads respectively) (2-way ANOVA, p-values = 0.8798 (DA) ; 0.6893 (AVJ) ; 0.9208 (CV) ; 0.2827 (ISVs) ; 0.7351 (CPCs)). (E) Injection of tumor cell lines with different cell diameters in the zebrafish embryo. (D) Ratios of cell diameter normalized to 1675 cell line and ratios of arrest occurring in small vessels (ISVs + CPCs) of the ZF embryo caudal plexus for 1675, D2A1, NIH3T3, A431, 4T1 and ZMEL-1 tumor cell lines. (Measure of diameter: n (cells) = 90 for all cell lines. Ratio of arrest in small vessels: n (embryos) = 28, 36, 27, 26, 30 and 20 for 1675, D2A1, NIH3T3, A431 and ZMEL-1 cell lines respectively.) (E) Correlation test of ratios of cell diameter normalized to 1675 cell line and ratios of arrest occurring in small vessels (D). (Measure of diameter: n (cells) = 90 for all cell lines. Ratio of arrest in small vessels: n (embryos) = 28, 36, 27, 26, 30 and 20 for 1675, D2A1, NIH3T3, A431 and ZMEL-1 cell lines respectively.) (Spearman r, p-value = 0.0028).

**Figure S2 related to Figure 2:**
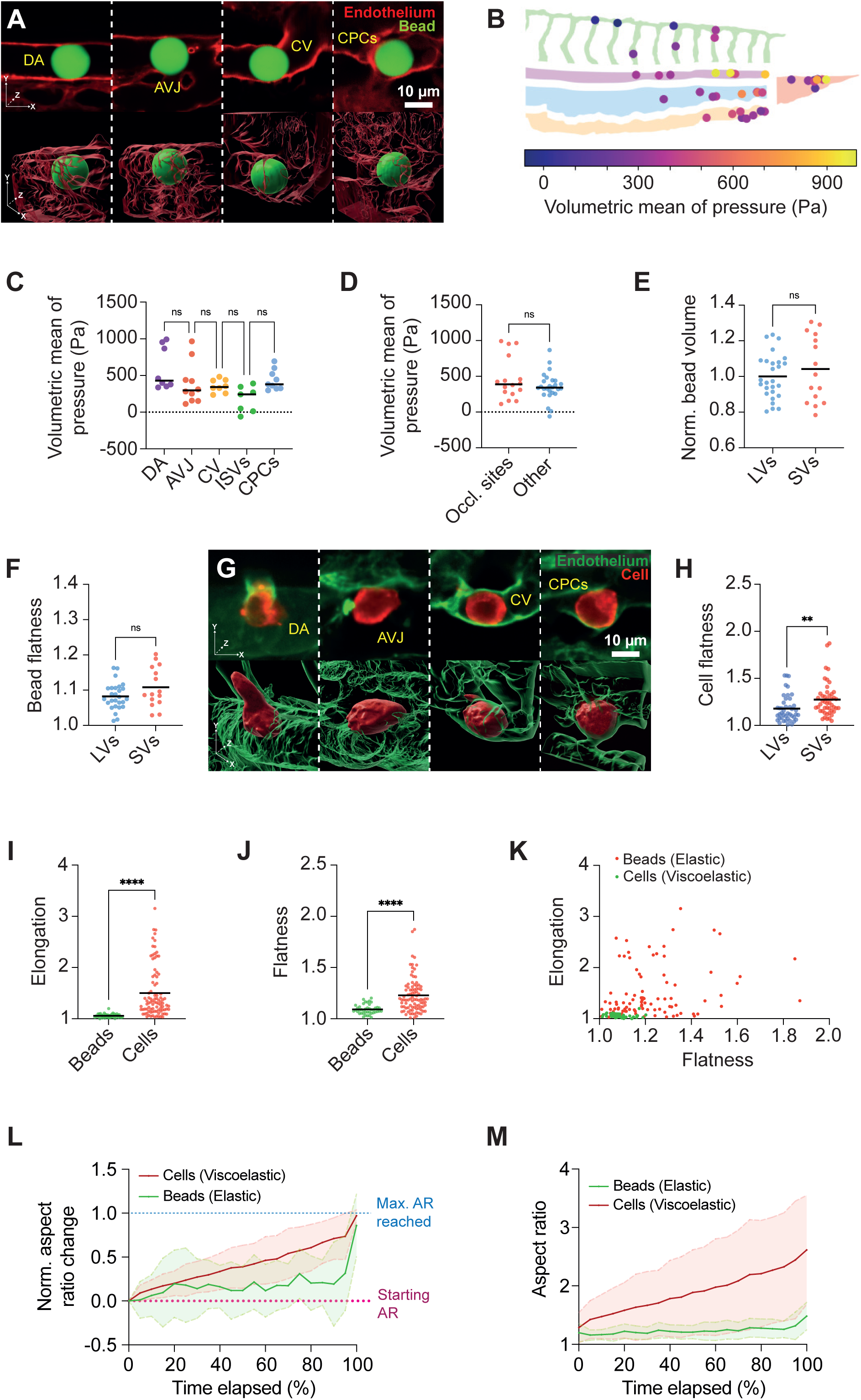
CTCs use viscous deformation to enter small-sized vessels. (A) Z-projection of confocal stack of arrested “D2A1-like” bead in the DA, AVJ, CV and CPCs. (B) Location and volumetric mean of pressure of arrested “D2A1-like” beads imaged and analyzed in the caudal plexus of the ZF embryo. (N = 6, n (beads) = 42). (C) Quantification of volumetric mean of pressure sustained by arrested beads in the different vascular subregions of the caudal plexus of the ZF embryo. (N = 6, n (beads) = 8, 10, 8, 7 and 9 for the DA, the AVJ, the CV, the ISVs and the CPCs respectively) (Kruskal-Wallis, p-values = 0.4044 (DA vs AVJ) ; > 0.9999 (AVJ vs CV) ; 0.8727 (CV vs ISVs) ; 0.1144 (ISVs vs CPCs)). (D) Quantification of volumetric mean of pressure sustained by arrested beads at occlusion sites vs all other sites. (N = 6, n (beads) = 16 and 26 for occlusion sites and other sites respectively) (Mann-Whitney, p-value = 0.4492). (E) Quantification of volume of beads arrested in LVs and SVs. (N = 6, n (beads) = 26 and 16 for LVs and SVs respectively) (Mann-Whitney, p-value = 0.4504). (F) Quantification of flatness of beads arrested in LVs and SVs. (N = 6, n (beads) = 26 and 16 for LVs and SVs respectively) (Mann-Whitney, p-value = 0.1283). (G) Z-projection of confocal stack of D2A1 tumor cells in the DA, AVJ, CV and CPCs. (H) Quantification of flatness of D2A1 tumor cells arrested in LVs and SVs. (N = 2, n (cells) = 44 and 47 for LVs and SVs respectively) (Mann-Whitney, p-value = 0.0017). (I) Quantification of elongation of beads and D2A1 tumor cells arrested in the caudal plexus of the ZF embryo. (N = 6 and 2 for beads and cells respectively, n (beads or cells) = 42 and 91 for beads and cells respectively) (Mann-Whitney, p-value = < 0.0001). (J) Quantification of flatness of beads and D2A1 tumor cells arrested in the caudal plexus of the ZF embryo. (N = 6 and 2 for beads and cells respectively, n (beads or cells) = 42 and 91 for beads and cells respectively) (Mann-Whitney, p-value = < 0.0001). (K) Plot of elongation and flatness factors of beads and D2A1 tumor cells arrested in the caudal plexus of the ZF embryo. (N = 6 and 2 for beads and cells respectively, n (beads or cells) = 42 and 91 for beads and cells respectively). (L) Normalized change in aspect ratio as a function of normalized time spent at ISV entrance for polyacrylamide elastic beads and D2A1 tumor cells arrested there for at least 10 seconds (N = 6 and 2 for beads and cells respectively, n (events of arrival, arrest of > 10 seconds and entry at DA-ISV connections) = 17 and 32 for beads and cells respectively). (M) Aspect ratio as a function of normalized time spent at ISV entrance for polyacrylamide elastic beads and D2A1 tumor cells arrested there for at least 10 seconds (N = 6 and 2 for beads and cells respectively, n (events of arrival, arrest of > 10 seconds and entry at DA-ISV connections) = 17 and 32 for beads and cells respectively).

**Figure S3 related to Figure 3:**
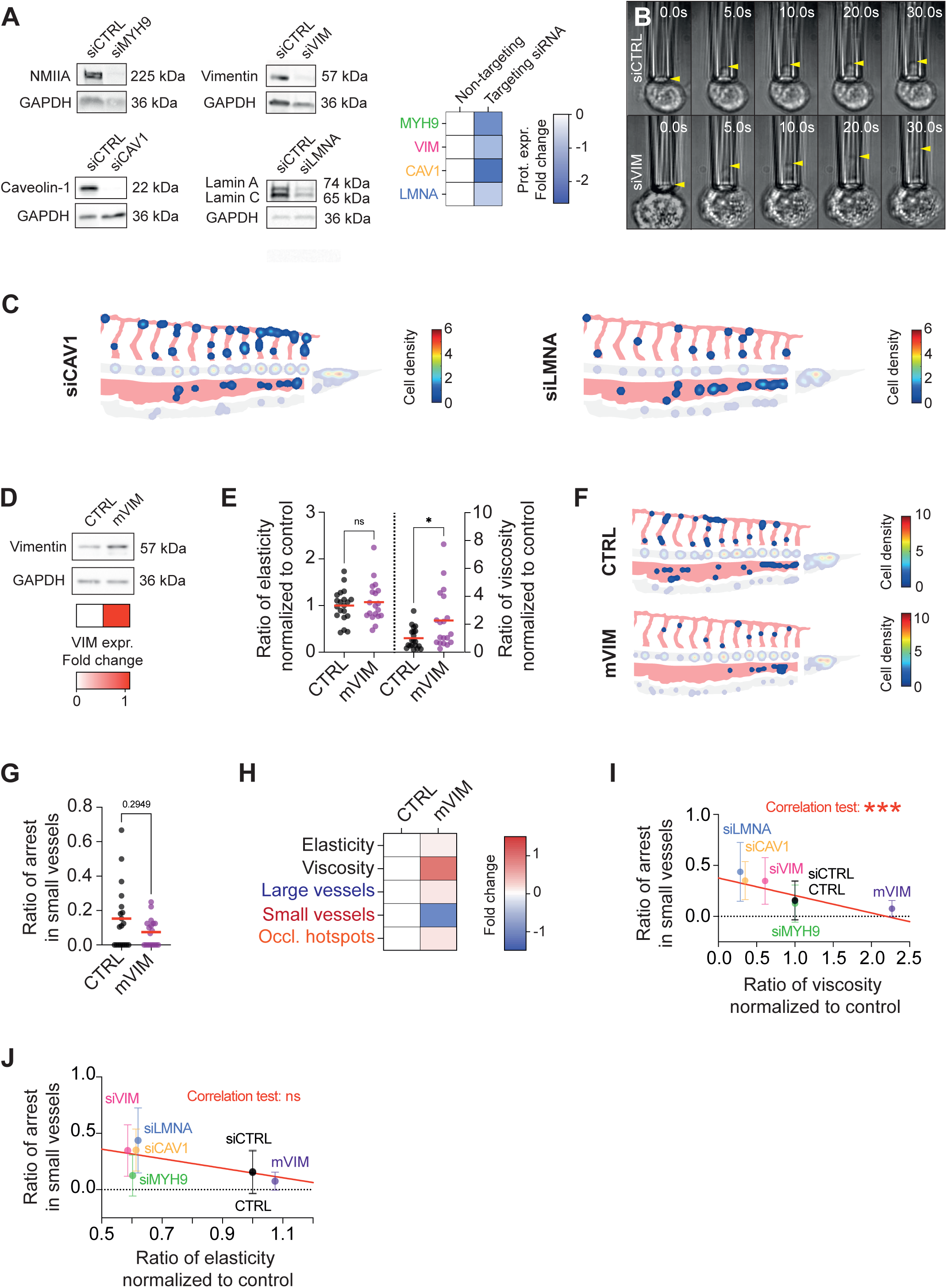
Cell viscosity dictates arrest locations of circulating tumor cells. (A) Representative WB images of protein expression of target-proteins 3 days post-transfection with siRNA and resulting protein expression fold changes. (B) Micropipette aspiration assay time lapses of siRNA-treated D2A1 tumor cells. Yellow arrowheads indicate length of cell aspirated in the capillary. (C) Heatmaps displaying hotspots of arrest 5 minutes post-injection of siRNA-treated D2A1 tumor cells in the ZF embryo caudal plexus (N = 3 (siCAV1, siLMNA), n (embryos) = 18 (siCAV1) ; 15 (siLMNA)). (D) Representative WB image of vimentin expression in mouse-vimentin transfected tumor cells and resulting vimentin expression fold change. (E) Ratios of elasticity and viscosity normalized to control of CTRL and mVIM-transfected D2A1 tumor cells. (N = 2, n (cells) = 20 and 19 for CTRL and mVIM respectively) (Welch’s t test of ratio of elasticity, p-value = 0.5534) (Man-Whitney test of ratio of viscosity, p-value = 0.0192). (F) Heatmaps displaying hotspots of arrest 5 minutes post-injection of CTRL and mVIM-transfected D2A1 tumor cells in the ZF embryo caudal plexus. (N = 4, n (embryos) = 19 and 18 for CTRL and mVIM respectively). (G) Quantification of ratio of arrest occurring in small vessels (ISVs + CPCs) of the ZF embryo caudal plexus for CTRL and mVIM-transfected D2A1 tumor cells. (N = 4, n (embryos) = 19 and 18 for CTRL and mVIM respectively) (Mann-Whitney, p-value = 0.2949). (H) Fold changes of ratios of elasticity, viscosity, arrest in small and large vessels and arrest at occlusion hotspots for CTRL- or mVIM-transfected tumor cells. (I) Correlation test of ratio of arrest occurring in small vessels and ratio of viscosity. (Spearman r, p-value = 0.0008) (J) Correlation test of ratio of arrest occurring in small vessels and ratio of elasticity. (Spearman r, p-value = 0.3571)

**Figure S4 related to Figure 4:**
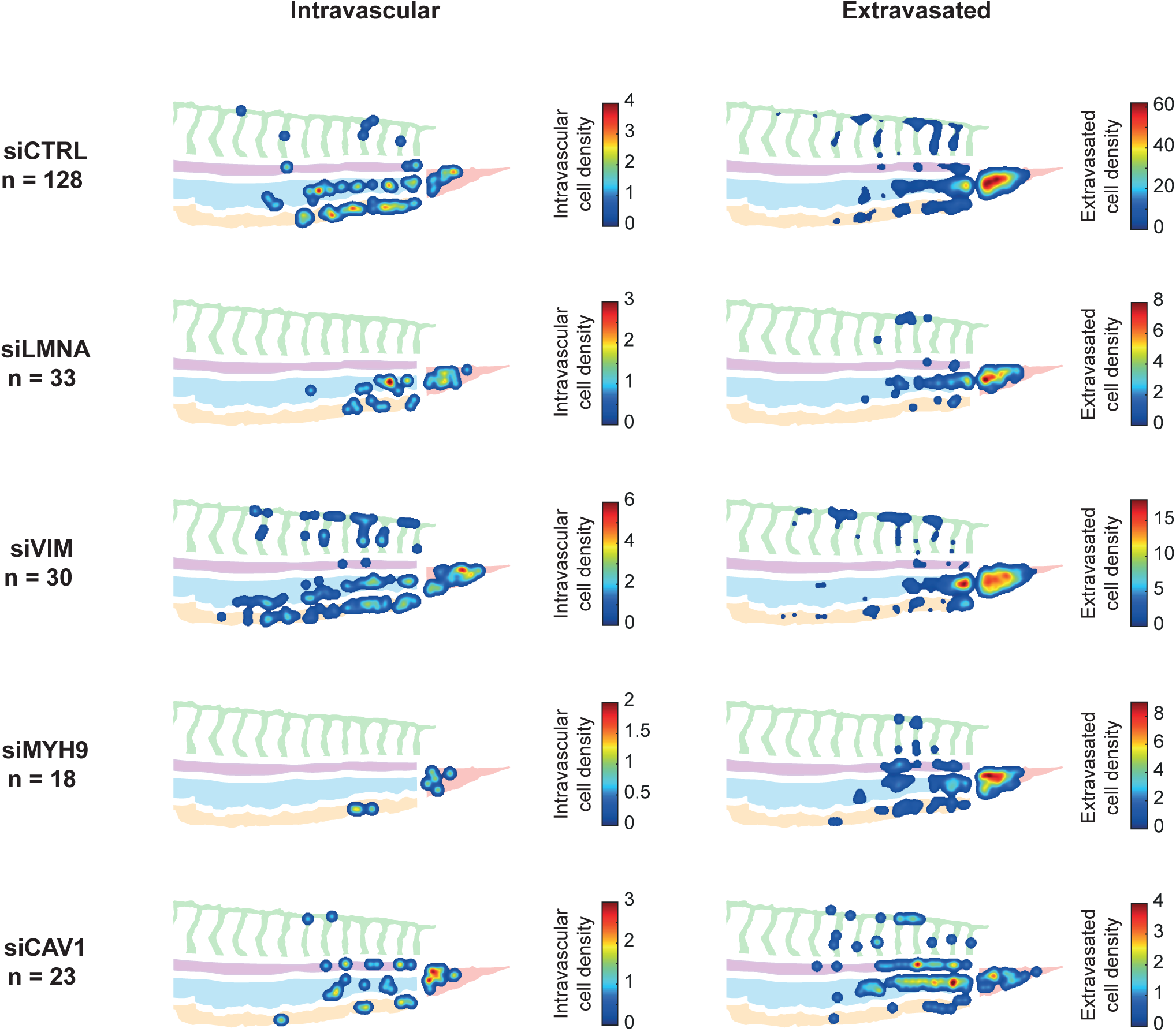
Tumor cell viscosity tunes metastatic extravasation. Heatmaps displaying hotspots of intravascular and extravasated siRNA-treated D2A1 tumor cells 24 hours post-injection. (N = 12 (siCTRL) ; 3 (siMYH9, siVIM, siCAV1, siLMNA), n (embryos) = 128 (siCTRL) ; 18 (siMYH9) ; 30 (siVIM) ; 23 (siCAV1) ; 33 (siLMNA)).

**Figure S5 related to Figure 4:**
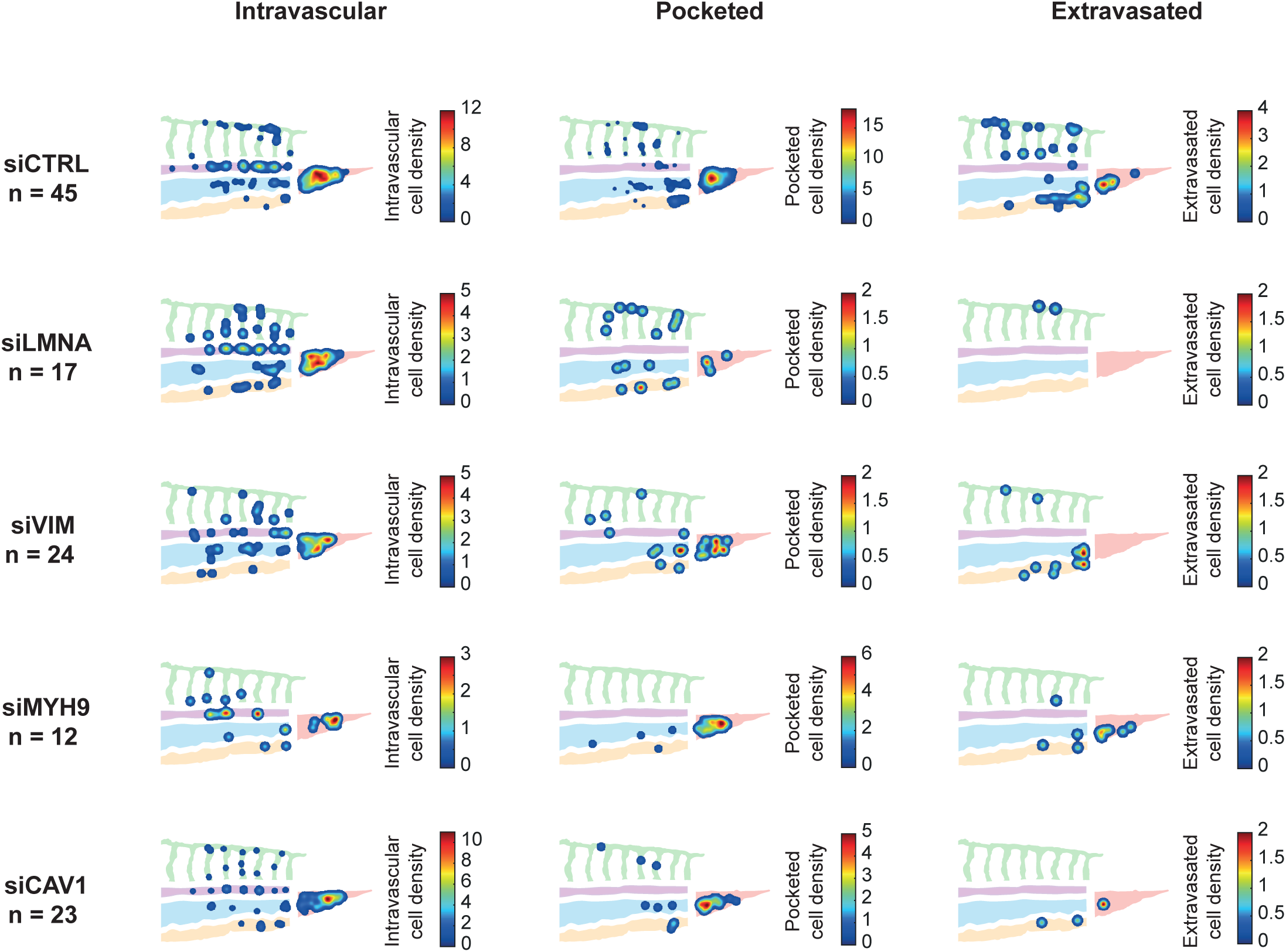
Tumor cell viscosity tunes metastatic extravasation. Heatmaps displaying hotspots of intravascular, pocketed and extravasated siRNA-treated D2A1 tumor cells 3 hours post-injection. (N = 4 (siCTRL) ; 2 (siMYH9, siVIM, siCAV1, siLMNA), n (embryos) = 45 (siCTRL) ; 12 (siMYH9) ; 24 (siVIM) ; 23 (siCAV1) ; 17 (siLMNA)).

**Figure S6 related to Figure 4:**
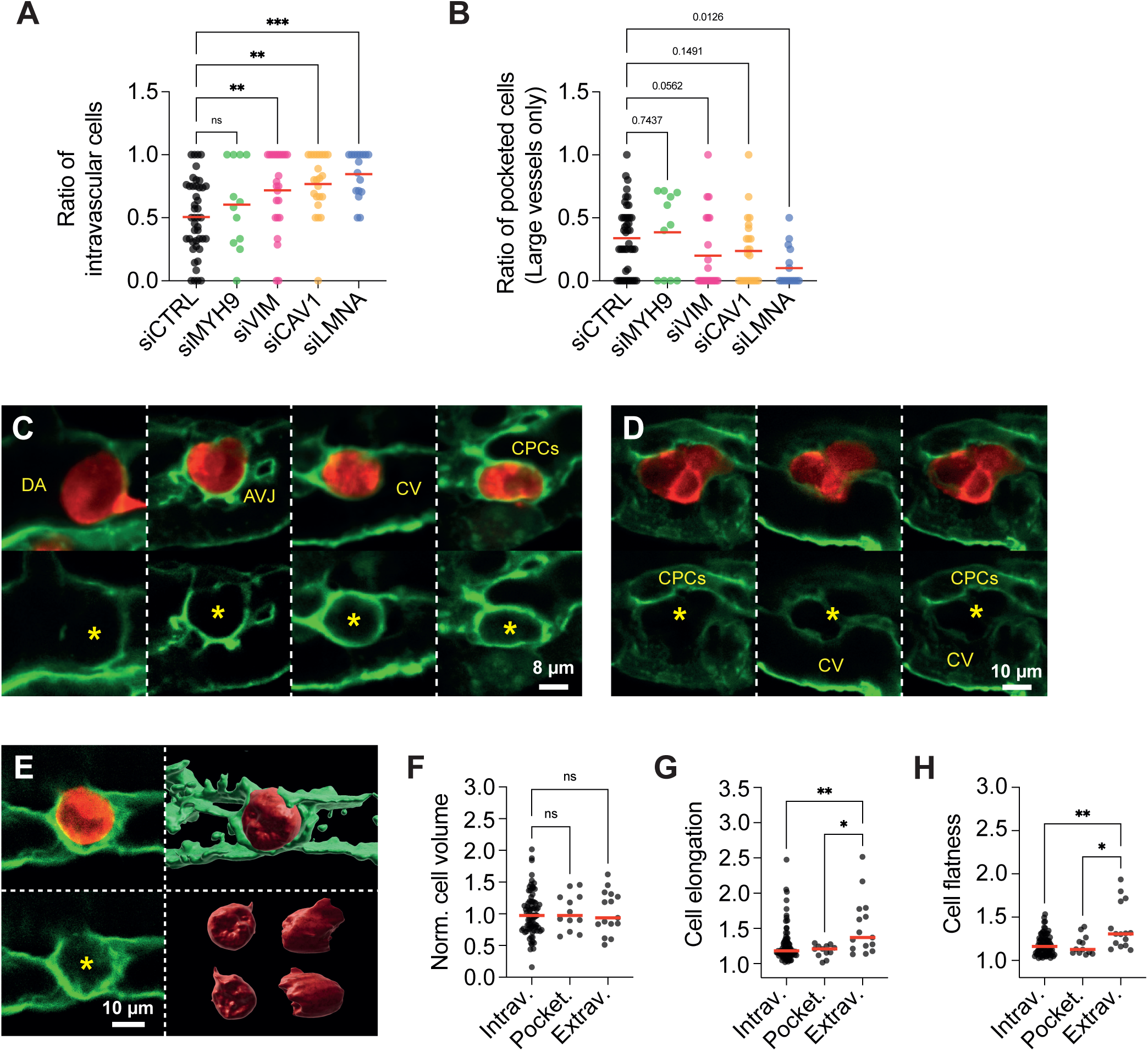
Tumor cell viscosity tunes metastatic extravasation. (A) Quantification of ratio of intravascular cells 3 hours post-injection of mechanically-altered tumor cells. (N = 4 (siCTRL) ; 2 (siMYH9, siVIM, siCAV1, siLMNA), n (embryos) = 45 (siCTRL) ; 12 (siMYH9) ; 24 (siVIM) ; 23 (siCAV1) ; 17 (siLMNA)) (Kruskal-Wallis, p-values = 0.3002 (siCTRL vs siMYH9) ; 0.0045 (siCTRL vs siVIM) ; 0.0014 (siCTRL vs siCAV1) ; 0.0004 (siCTRL vs siLMNA)). (B) Quantification of ratio of pocketed cells in large vessels of the ZF embryo caudal plexus 3 hours post-injection of mechanically-altered tumor cells. (N = 4 (siCTRL) ; 2 (siMYH9, siVIM, siCAV1, siLMNA), n (embryos) = 44 (siCTRL) ; 11 (siMYH9) ; 22 (siVIM) ; 23 (siCAV1) ; 16 (siLMNA)) (Kruskal-Wallis, p-values = 0.7437 (siCTRL vs siMYH9) ; 0.0562 (siCTRL vs siVIM) ; 0.1491 (siCTRL vs siCAV1) ; 0.0126 (siCTRL vs siLMNA)). (C) Z-projections of confocal stacks of pocketed D2A1 tumor cells in the DA, AVJ, CV and CPCs 3 hours post-injection. Yellow stars indicate tumor cell-containing endothelial pockets. (D) Z-projections of a single confocal stack of D2A1 tumor cells extravasated between the CV and the CPCs. Yellow stars indicate the location of the cells in the extravascular space between the CV and the CPCs. (E) Z-projection of confocal stack of pocketed D2A1 tumor cell 3 hours post-injection and corresponding 3D renders. Yellow star indicates tumor cell-containing endothelial pocket. (F) Quantification of normalized volume of intravascular, pocketed and extravasated D2A1 tumor cells 3 hours post-injection. (N = 2, n (cells) = 78, 12 and 15 for intravascular, pocketed and extravasated cells respectively) (Ordinary one-way ANOVA, p-values = 0.7696 (Intrav. vs Pocket. and Intrav. vs Extrav.) (G) Quantification of elongation of intravascular, pocketed and extravasated D2A1 tumor cells 3 hours post-injection. (N = 2, n (cells) = 78, 12 and 15 for intravascular, pocketed and extravasated cells respectively) (Kruskal-Wallis, p-values = 0.6183 (Intrav. vs Pocket.) ; 0.0067 (Intrav. vs Extrav.) ; 0.0130 (Pocket. vs Extrav.). (H) Quantification of flatness of intravascular, pocketed and extravasated D2A1 tumor cells 3 hours post-injection. (N = 2, n (cells) = 78, 12 and 15 for intravascular, pocketed and extravasated cells respectively) (Kruskal-Wallis, p-values = 0.9610 (Intrav. vs Pocket.) ; 0.0033 (Intrav. vs Extrav.) ; 0.0234 (Pocket. vs Extrav.).

**Figure S7 related to Figure 5:**
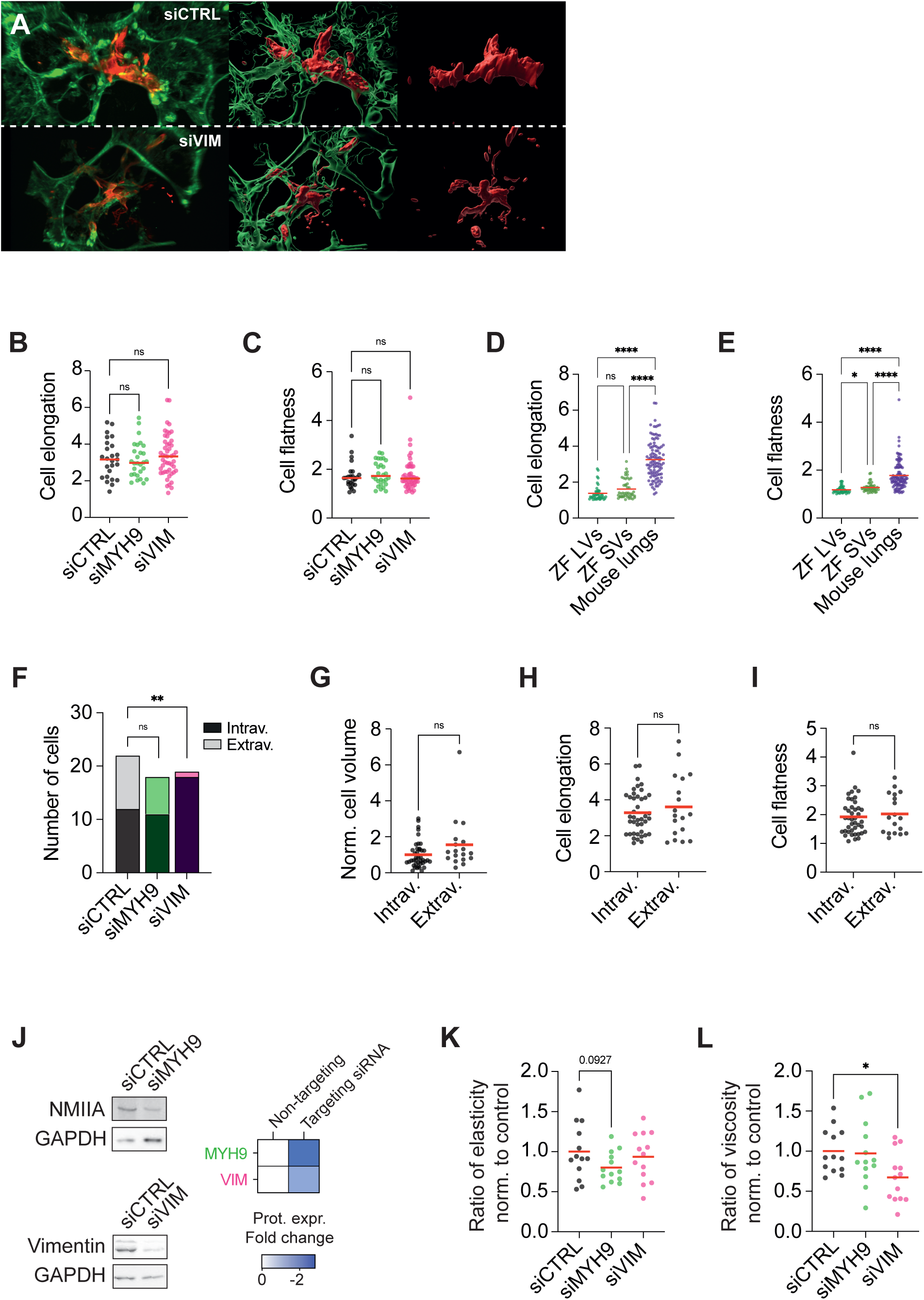
The viscoelastic profile of CTCs has a lasting impact on post-extravasation metastatic outgrowth. (A) Z-projections of confocal stacks of mechanically-altered D2A1 tumor cells arrested in the mouse lung vasculature 1-hour post-injection and corresponding 3D renders. (B) Quantification of elongation of mechanically-altered D2A1 tumor cells arrested in the mouse lung vasculature 1-hour post-injection (N = 1, n (cells) = 25 (siCTRL) ; 28 (siMYH9) ; 49 (siVIM)) (Kruskal-Wallis, p-values = 0.8081 (siCTRL vs siMYH9 and siCTRL vs siVIM)). (C) Quantification of flatness of mechanically-altered D2A1 tumor cells arrested in the mouse lung vasculature 1-hour post-injection (N = 1, n (cells) = 25 (siCTRL) ; 28 (siMYH9) ; 49 (siVIM)) (Kruskal-Wallis, p-values = 0.7983 (siCTRL vs MYH9) ; 0.9964 (siCTRL vs siVIM) (D) Comparison of cell elongation values found in the zebrafish embryo and mouse lung vasculatures. (N = 2 (ZF) ; 1 (Mouse), n (cells) = 44 (Zebrafish LVs) ; 47 (Zebrafish SVs) ; 102 (Mouse lungs)) (Kruskal-Wallis, p-values = 0.1263 (ZF LVs vs ZF SVs) ; < 0.0001 (ZF LVs vs Mouse Lungs) ; < 0.0001 (ZF SVs vs Mouse Lungs)). (E) Comparison of cell flatness values found in the zebrafish embryo and mouse lung vasculatures. (N = 2 (ZF) ; 1 (Mouse), n (cells) = 44 (Zebrafish LVs) ; 47 (Zebrafish SVs) ; 102 (Mouse lungs)) (Kruskal-Wallis, p-values = 0.0443 (ZF LVs vs ZF SVs) ; < 0.0001 (ZF LVs vs Mouse Lungs) ; < 0.0001 (ZF SVs vs Mouse Lungs)). (F) Quantification of intravascular and extravascular mechanically-altered D2A1 tumor cells in the mouse lung vasculature 24 hours post-injection. (N = 2, n (cells) = 22 (siCTRL) ; 18 (siMYH9) ; 19 (siVIM)) (Fischer’s exact test performed with raw number of cells, p-values = 0.7547 (siCTRL vs siMYH9) ; 0.0048 (siCTRL vs siVIM)). (G) Quantification of volume normalized to control of intravascular and extravasated D2A1 tumor cells in the mouse lung vasculature 24 hours post-injection. (N = 2, n = 41 and 18 for intravascular and extravasated cells respectively) (Mann-Whitney, p-value = 0.0707). (H) Quantification of elongation of intravascular and extravasated D2A1 tumor cells in the mouse lung vasculature 24 hours post-injection (N = 2, n = 41 and 18 for intravascular and extravasated cells respectively) (Mann-Whitney, p-value = 0.6537). (I) Quantification of flatness of intravascular and extravasated D2A1 tumor cells in the mouse lung vasculature 24 hours post-injection (N = 2, n = 41 and 18 for intravascular and extravasated cells respectively) (Mann-Whitney, p-value = 0.6774). (J) Representative WB images of protein expression of target-proteins 7 days post-transfection with siRNA and resulting protein expression fold changes. (K) Ratio of elasticity of siRNA-treated D2A1 tumor cells 7 days post-transfection normalized to control. (N = 1, n = 13 for siCTRL, siVIM and siMYH9) (Ordinary one-way ANOVA, p-values = 0.5838 (siCTRL vs siVIM) ; 0.0927 (siCTRL vs siMYH9)). (L) Ratio of viscosity of siRNA-treated D2A1 tumor cells 7 days post-transfection normalized to control. (N = 1, n = 13 for siCTRL, siVIM and siMYH9) (Ordinary one-way ANOVA, p-values = 0.0187 (siCTRL vs siVIM) ; 0.8358 (siCTRL vs siMYH9)).

